# Temporal profiling of human lymphoid tissues reveals coordinated defence to viral challenge

**DOI:** 10.1101/2023.09.15.558006

**Authors:** Matthew L. Coates, Nathan Richoz, Zewen K. Tuong, Georgie Bowyer, Colin Y.C. Lee, John R. Ferdinand, Eleanor Gillman, Mark McClure, Rafael Di Marco Barros, Benjamin J Stewart, Menna R. Clatworthy

## Abstract

Adaptive immunity is generated in lymphoid organs, but how these structures defend themselves during infection in humans is unknown. The nasal epithelium is a major site of viral entry, with adenoid nasal-associated lymphoid tissue (NALT) generating early adaptive responses. Here, using a nasopharyngeal biopsy, we examined longitudinal immune responses in NALT following viral challenge, using SARS-CoV-2 infection as a natural experimental model. In acute infection, infiltrating monocytes formed a subepithelial and peri-follicular shield, recruiting NET-forming neutrophils, whilst tissue macrophages expressed pro-repair molecules during convalescence to promote the restoration of tissue integrity. Germinal centre B cells expressed anti-viral transcripts that inversely correlated with fate-defining transcription factors. Among T cells, tissue-resident memory CD8 T cells alone showed clonal expansion and maintained cytotoxic transcriptional programmes into convalescence. Together our study provides a unique insight into how human nasal adaptive immune responses are generated and sustained in the face of viral challenge.

---

Humoral adaptive immune responses generate high affinity antibodies that are important for defence against pathogenic immune challenge. This process is initiated when B cell receptor (BCR)-bound antigen is internalised, processed and presented in the context of surface major histocompatibility complex (MHC) class II to a cognate CD4 T cell, initiating a germinal centre (GC) response which typically occurs in secondary lymphoid organs ^1^. GCs are highly organised structures with distinct light and dark zones. Germinal centre light zones contain not only GC B cells, but also follicular dendritic cells (FDC), and PD1+ CD4+ T follicular helper (Tfh) cells. The interaction of these three cell types leads to iterative rounds of GC B cell proliferation, during which there is somatic hypermutation and class switching from IgM to other isotypes, with the emergence of both memory B cells and antibody secreting plasmablasts/cells that defend against current and future infection^2–7^. As an important site in the generation of long-term immunity, lymphoid tissues need a robust defence mechanism of their own. In mice, functionally diverse macrophage populations such as sub-capsular sinus macrophages, co-ordinate lymph node defence by innate lymphocyte crosstalk that prevents the spread of lymph-borne viruses and bacterial infections ^8,9^.

These cell type-specific molecular processes underpinning the generation and defence of adaptive immune responses in secondary lymphoid organs, such as lymph node and spleen, are well studied in animal models. In contrast, functional profiling of human lymphoid tissue responses is limited, with stimulation of representative adaptive immune responses and collection of appropriate tissue samples constituting major barriers. The nasopharynx contains the pharyngeal tonsils (also known as adenoids) that form part of Waldeyer’s ring of lymphoid tissue. Pharyngeal tonsils are the major site of nasal-associated lymphoid tissue (NALT) and have a surface crypt epithelium with antigen-transporting M-cells and underlying B cell follicles and T cell-containing interfollicular areas, that together enable the generation of protective antibody-secreting and memory cells following nasal challenge ^10,11^. CD68+ macrophages have also been identified in NALT ^12^, but cell subset heterogeneity, and their contribution to tissue defence or repair post-infection is completely unknown in humans.

Since NALT can be readily accessed in humans using an endoscope, we investigated whether NALT biopsies may offer a feasible method to profile early adaptive immune responses in secondary lymphoid organs following an upper respiratory tract immune challenge. Furthermore, we sought to understand how these tissues defend themselves in the face of local infection, to ensure that the cells and infrastructure required to support an ongoing immune response are maintained. We developed an endoscopic technique for collecting NALT biopsies from live human subjects using topical local anaesthesia. We used this methodology to undertake a ‘natural experiment’, sampling adult subjects at two stages of SARS-CoV-2 infection – a virus that uses nasal epithelium as a point of entry and replication^13,14^.

Although SARS-CoV-2 is already well studied in human subjects, much of this work has used peripheral blood or epithelial surface sampling in live subjects, and thus fail to capture and profile lymphoid follicular components, such as germinal centre B cells or bona-fide T follicular helper cell populations^15–17^. Lymphoid tissue sampling in SARS-COV-2 infection has been conducted in children suggesting a robust germinal centre response in NALT whilst, in contrast, autopsy samples from spleen and lymph nodes in adults with fatal SARS-CoV-2 infection suggest a disruption of the germinal centre process in fatally severe disease^18–20^. We applied single-cell RNA sequencing (scRNAseq), multi-parameter flow cytometry and confocal imaging to NALT biopsies, to understand the cellular molecular adaptations that support, polarize and defend human secondary lymphoid tissue in adults in the face of a viral immune challenge. (**Fig. 1A, S1B**)

**Figure 1.**
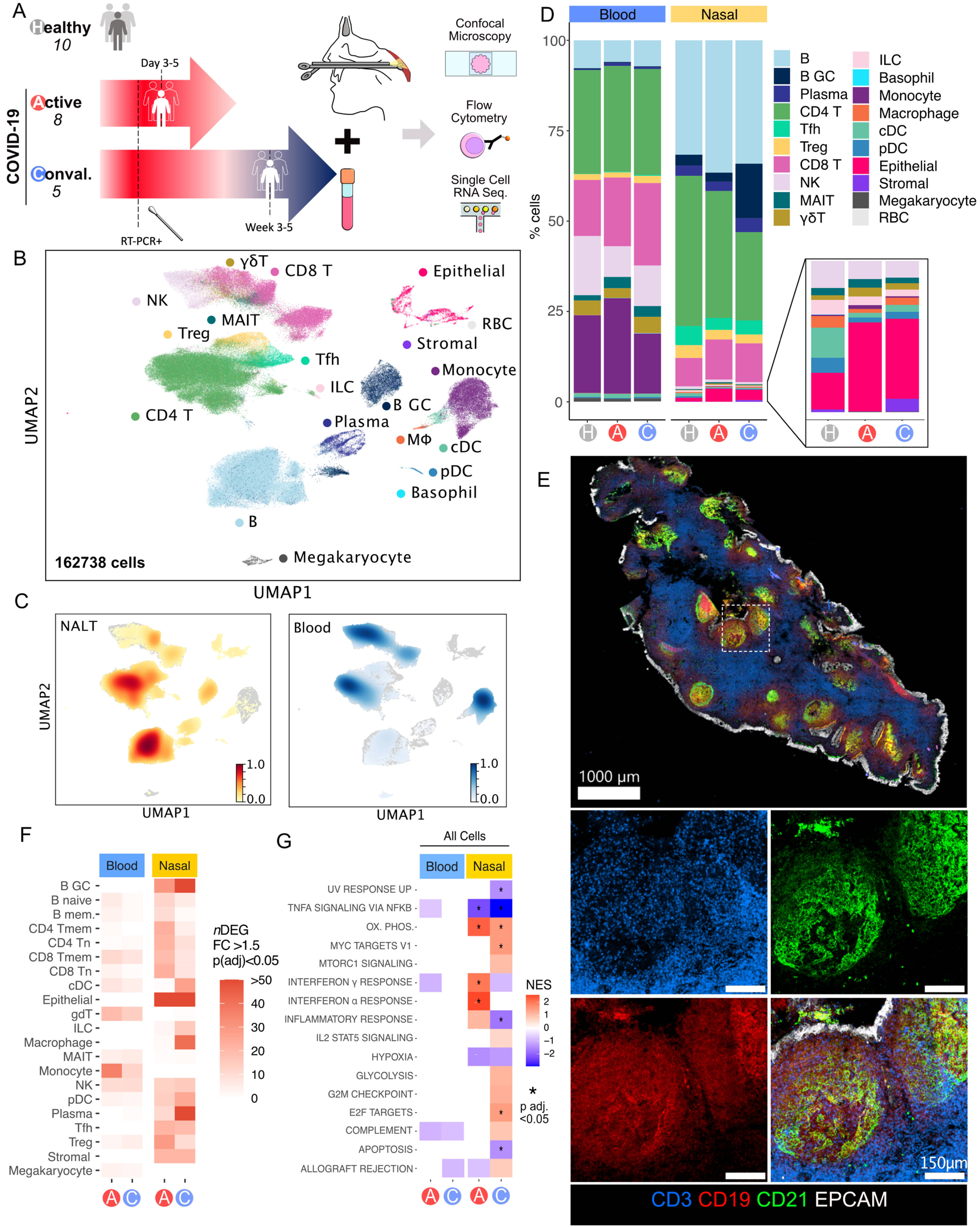
Experiment overview and cellular landscape of blood and NALT in SARS CoV-2 infection. **(A)** Schematic of experimental design, showing groups, numbers of subjects per group, and sampling strategy. Conval., Convalescent **(B)** Uniform manifold approximation and projection (UMAP) of all cells in COVID-19 subjects and healthy controls, labelled with major cell annotations. B GC, Germinal centre B cell; ILC, Innate Lymphoid Cell; MAIT, Mucosal-associated invariant T cell; MΦ, Macrophage; NK, Natural Killer cell; Tfh, T follicular helper cell; Treg, T regulatory cell; cDC, conventional dendritic cell; γδT, Gamma-Delta T cell; pDC, Plasmacytoid dendritic cell; RBC, Red blood cell **(C)** UMAP of embedding density (scaled Gaussian kernel density estimation) of all cells by sample type. Nasal associated lymphoid tissue (NALT) embedding density is shown in orange, and Blood embedding density is shown in blue. **(D)** Stacked bar charts showing proportional split (as percentage) of NALT and blood samples by cell type and disease type, with nested bar chart showing proportions, by disease type, of nasal NK, MAIT, γδT, ILC, monocyte, macrophage, cDC, pDC, Epithelial and Stromal cells. A, Active COVID; C, Convalescent COVID; H, Healthy control. **(E)** Confocal immunofluorescence microscopy of section of a postnasal space biopsy sample in 19 year old female from the Convalescent COVID-19 group, showing presence of lymphoid follicles. White dashed box indicates region magnified in the 4 boxes in the lower part of the image panel. Red, CD19; Green, CD21; White, EpCAM; Blue, CD3. **(F)** Number of differentially expressed genes (*n*DEG) for disease vs. healthy control samples in NALT and Blood samples by cell type cluster and disease type, where expression Fold Change (FC) >1.5 and Bonferroni adjusted p value (p(adj)) <0.05. A full list of abbreviations can be found in the Glossary of Terms. **(G)** Hallmark Gene set enrichment analysis of differentially expressed genes between disease and healthy control subjects for all (combined) cells by sample type and disease state. NES, Normalised enrichment score: *, Benjamini-Hochberg adjusted p value (by permutation) <0.05

## Results

### Nasal biopsy enables profiling of cellular profiling of human mucosal-associated lymphoid tissue

23 subjects were sampled (aged 19-91 years), including 10 healthy controls, 8 acute COVID-19 patients (sampled within one week of a first positive SARS-CoV-2 RT-PCR test), and 5 convalescent COVID-19 patients (clinically asymptomatic subjects sampled 3-5 weeks post-first positive SARS-CoV-2 RT-PCR, **Fig. 1A**, **S1A-B**). Post-nasal space biopsies (with additional nasal brushing/curettage in some cases) were collected, alongside paired blood samples, and processed for flow cytometry, scRNAseq, and confocal microscopy (**Fig. S1A**).

Post-QC, we generated data on 162,738 cells and identified 20 major cell clusters across peripheral blood mononuclear cells (PBMC) and nasal tissues, including epithelial cells, stromal cells and a variety of immune cell populations (**Fig. 1B-D, S1C**). While most cell subsets could be found in both NALT and blood, there was a significant over-representation of GC B cells and Tfh cells in nasal samples (**Fig. 1C-D**), reflecting the presence of GC-containing lymphoid structures. GC B cells were particularly abundant in convalescent NALT samples, consistent with the expansion of virus-specific responses (**Fig. 1D)**. NALT samples also differ from adult nasal and bronchial brushings^17^, in which Tfh and GC B cells are scarce **(Fig. S1D-F)**. Confocal imaging of NALT biopsies confirmed the presence of organised lymphoid structures, including B cell follicles containing CD21-expressing FDCs and spatially distinct T cell-rich areas, with overlying surface EPCAM+ epithelial cells (**Fig. 1E**). Notably, the number of differentially expressed genes (DEGs) in COVID-19 samples, (both acute (A) and convalescent (C)), relative to controls, was greater in NALT than in blood (**Fig. 1F**), emphasising the utility of this tissue sampling strategy to deliver improved insights into immune responses in COVID-19. Similarly, gene set enrichment analysis (GSEA) of these DEGs showed a greater magnitude, significance and number of enriched gene-sets in NALT compared with peripheral blood, including *Interferon alpha response* pathway genes **(Fig 1G)**.

Overall, these data show that nasopharyngeal biopsies, can be used to profile functional lymphoid tissue in adults, with capture of distinct follicular cell types not found in blood or nasal cavity brushings. NALT sampling presents a feasible method to interrogate the cellular molecular processing occurring during the generation of adaptive immune responses to a nasal viral infection in humans.

### Distinct temporal variation in the contribution of different MNP subsets to lymphoid tissue defence and repair

Post-nasal space sampling 3-5 days after diagnostic testing for SARS-CoV-2 provided a unique opportunity to understand how early innate immune responses act to defend NALT from local viral attack. Among mononuclear phagocytes (MNPs), we identified 11 cell types using canonical marker gene expression: conventional dendritic cells (cDC) (*CD141*^+^ cDC1 and *CD1C*^+^ cDC2), plasmacytoid DCs (pDC), six monocyte clusters (classical, intermediate, non-classical, *SIGLEC1*^+^, *CD163*^hi^, *C1Q*^+^), basophils and macrophages (**Fig. 2A**, **S2A**). Notably, the proportion of classical monocytes in nasal tissue increased in acute COVID-19 compared to healthy control and convalescent COVID-19 (**Fig. 2B-C**), consistent with tissue infiltration (confirmed by flow cytometry (**Fig. S2B**)), and these monocytes also showed increased expression of several self- and neutrophil-recruiting chemokines (*CCL2* and *CXCL2*/*CXCL3*/*CXCL8* respectively) (**Fig. 2D**). S100A8/9 transcripts were also increased in classical monocytes in active and convalescent COVID-19 samples (**Fig. 2D-E, S2C**). S100A8/9 dimerises to form calprotectin, which sequesters metal ions limiting their availability to bacteria, including upper respiratory tract-dwelling organisms such as Staphylococci ^21^. SARS-CoV-2 is known to directly infect respiratory tract epithelium, and indeed, we found SARS-CoV-2 Spike protein positive epithelial cells overlying NALT in acute COVID-19 (**Fig. 2F, S2D**). In this context, CD14+ monocytes were located in subepithelial regions of NALT in acute COVID-19 and surrounding some B cell follicles in convalescent disease (**Fig. 2G**), forming a shield around the structures required to generate early mucosal humoral immune responses. In addition, neutrophils were also present in the epithelial and subepithelial areas, some with histone citrullination, required for formation of neutrophil extracellular traps (NET) - structures that bind and kill invading bacteria ^22^ (**Fig. S2E**). Therefore, calprotectin-producing monocytes and NET-osing neutrophils may work together to defend the underlying lymphoid tissues following epithelial cell damage by invading virus.

**Figure 2.**
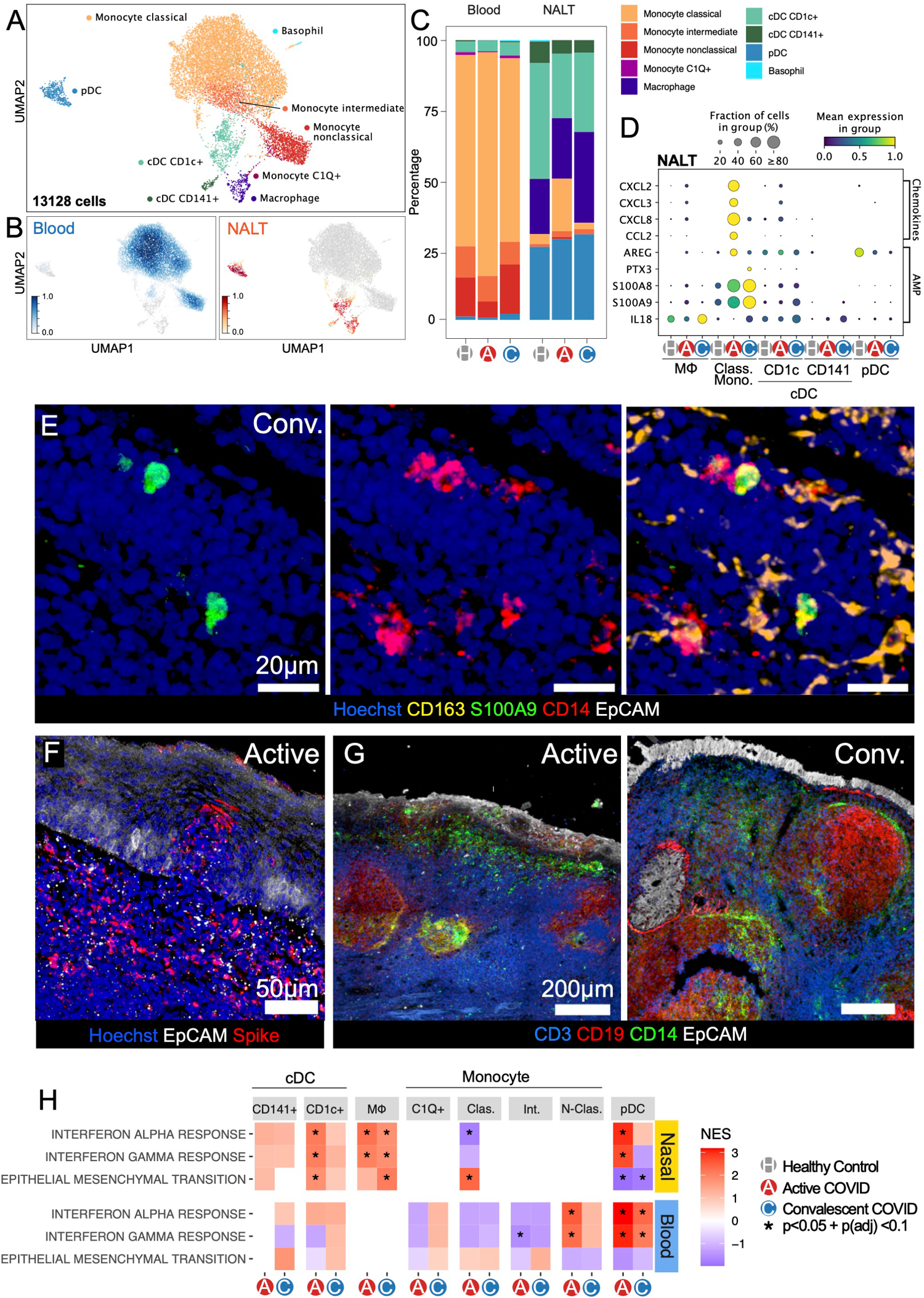
Recruited monocytes and inflammatory DCs form defensive shield in sub-epithelium. **(A)** Uniform manifold approximation and projection (UMAP) of mononuclear phagocyte (MNP) labelled cells, subset and re-clustered in isolation, with assigned subset cell type cluster labels shown. **(B)** Embedding density (Gaussian kernel estimation) UMAP of re-clustered MNP subset cell types showing scaled density of cells by sample type (Blood/ Nasal) where 1 = High density. NALT, orange; blood, blue. **(C)** Stacked bar chart showing proportional representation of MNP cell types in blood and nasal samples by disease status. A, Active COVID-19, C, Convalescent COVID-19, H, Healthy control **(D)** Selected chemokine and antimicrobial peptide (AMP) scaled expression in NALT MNP subsets. cDC, Conventional Dendritic Cell; Class. Mono., Classical monocyte; MΦ, Macrophage; pDC, Plasmacytoid dendritic cell. Size of dot indicates fraction of cells in group expressing gene, and colour indicates scaled mean gene expression. **(E)** Immunofluorescence microscopy of NALT from a convalescent COVID-19 subject, showing co-expression of CD14 and S100A9 in NALT tissue. Blue, Hoechst, Yellow, CD163, Green, S100A9, Red, CD14, White, EpCAM **(F)** Immunofluorescence microscopy showing expression of SARS-COV-2 spike protein in NALT epithelium from an active COVID-19 subject. Hoechst, Blue; SARS-COV-2 spike protein, Red; EpCAM, White. **(G)** Immunofluorescence microscopy comparing spatial localisation of CD14+ cells in NALT from subjects with active COVID-19 infection (left) and Convalescent COVID-19 infection (right). CD3, Blue; CD19, Red; CD14, Green; EpCAM, White. **(H)** Heatmap of GSEA Hallmark gene-set enrichment for differentially expressed genes in MNP subsets by disease group against healthy control samples. Colour of tile shows normalised enrichment score (NES). Red, Increased gene-set enrichment relative to healthy controls; Blue, Decreased gene-set enrichment relative to healthy controls; white/no colour, no enrichment of gene-set), * Statistical significance (p-value<0.05 and Benjamini-Hochberg adjusted p-value <0.1).

Macrophages were found exclusively in nasal samples, and showed delayed expansion relative to monocytes, increasing in convalescent COVID-19. The number of DEG in NALT-resident macrophages was also greater in convalescent COVID-19 compared with acute infection (**Fig. 1F**), including IL18 (**Fig. 2D**), reminiscent of studies in murine lymph nodes showing macrophage production of IL18 in the context of bacterial challenge ^9^. Interestingly, a population of circulating *C1Q*+ monocytes were also expanded in convalescent COVID-19 (**Fig 2C, S2A)**, with trajectory analysis suggesting that these represent tissue macrophage precursors (**Fig S2F)**.

Gene set enrichment analysis (GSEA) ^23^ showed a greater magnitude of enrichment in nasal cell types compared with their circulating counterparts (**Fig. 2H, S2G**). Considering the majority myeloid populations in NALT, *‘Interferon alpha response*’ and ‘*Interferon gamma response*’ genes were significantly enriched in macrophages, *CD1C*+ DC and pDC in acute COVID-19 compared to controls (**Fig. 2H, S2G**). In convalescent COVID-19, this interferon response gene enrichment became less prominent in cDC and pDC but was sustained in macrophages (**Fig. 2H**). Furthermore, ‘*epithelial-mesenchymal transition*’ gene-set enrichment increased in macrophages in convalescent disease, and indeed, was exclusively enriched in macrophages at this time-point (**Fig. 2H, S2G**), consistent with a role in the restoration of tissue homeostasis.

To further explore myeloid cell heterogeneity in NALT, we re-clustered nasal MNPs in isolation from those in blood (**Fig 3A**). This enabled the further identification of so-called ‘inflammatory’ or ‘monocyte-derived’ DCs (moDC, based on the expression of *CD14*, *S100A8/9* and *FCGR1A*(CD64) ^24,25^, and two distinct clusters of macrophages based on *C1Q* and *FOLR2* expression (**Fig. 3A, S3A**). The *FOLR2*+ macrophages expressed additional markers associated with tissue-resident macrophages, including *TIMD4* (TIM4) ^26^ (**Fig S3A**).

**Figure 3.**
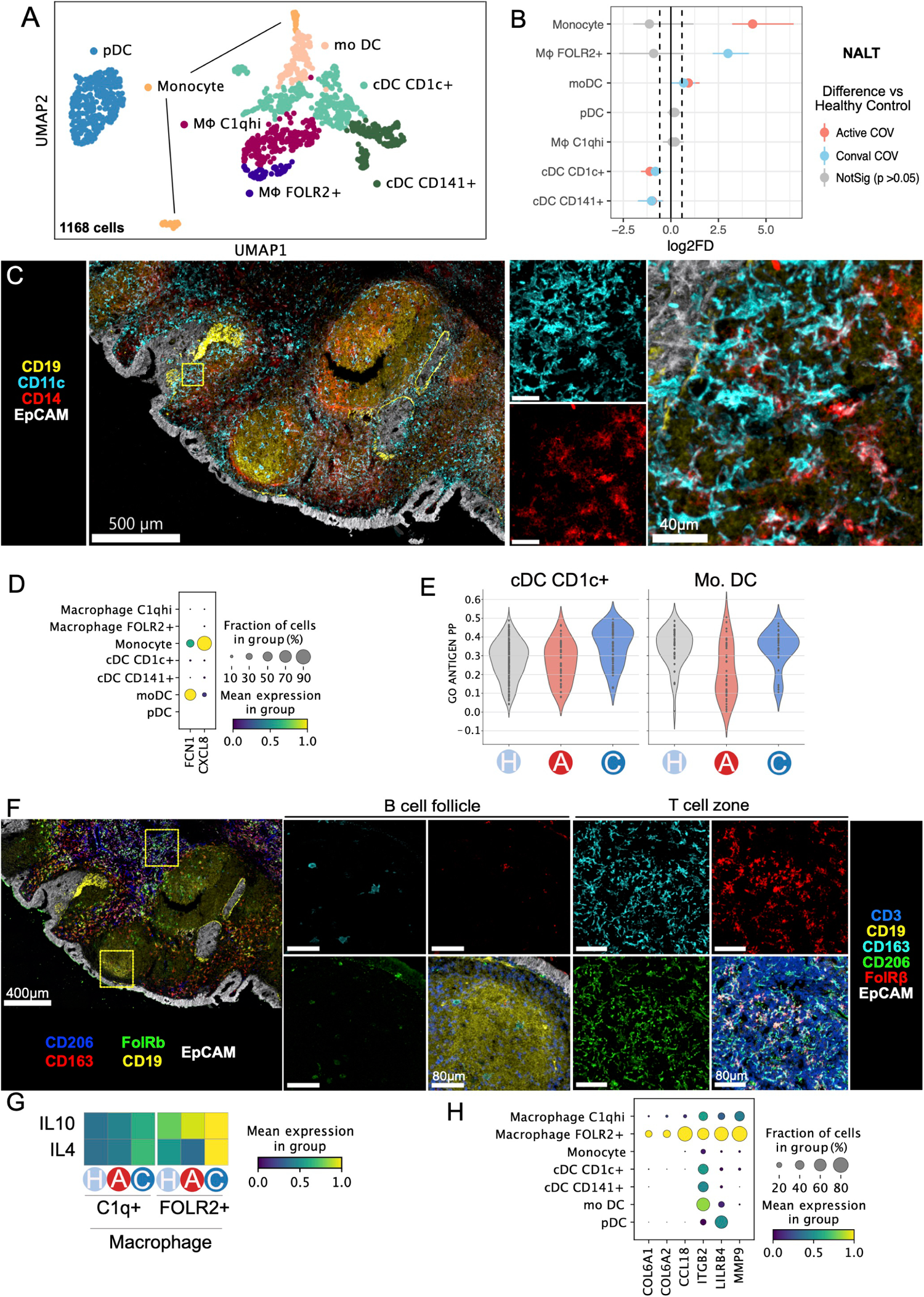
Tissue resident macrophages take on pro-repair phenotype in convalescence. **(A)**UMAP of re-clustered NALT MNP cell types in isolation. **(B)**Proportional difference in abundance of NALT MNP cell types in Active COVID-19 (red) and Convalescent COVID-19 (blue), relative to healthy controls. Non-significant values (p>0.05) relative to healthy controls, following Permutation testing, are shown in grey. Bars indicate bootstrapped 95% confidence interval. **(C)** Confocal microscopy imaging of NALT in Convalescent COVID-19 subject showing cellular co-expression of CD14+ (red) and CD11c+ (cyan), and their spatial localisation within NALT tissue relative to B cell follicles (CD19, indicated in yellow) and epithelium (EpCAM, indicated in white). Yellow box indicates area of increased magnification seen in boxes to the right of the image.**(D)** Dotplot showing expression of *FCN1* and *CXCL8* in NALT MNP subsets. Size of point indicates fraction of cells in each group expressing gene, colour of point indicates scaled mean expression of gene in each cell type group. **(E)** Violin plot showing expression of antigen processing and presentation Gene Ontology term genes (GO Antigen PP) in conventional CD1c+ cDC compared with Monocyte-derived DC (MoDC) cell subsets. Each point represents a cell. Healthy controls (H), Active COVID-19 (A), Convalescent COVID-19 (C). **(F)** Confocal imaging of NALT in convalescence showing distribution of macrophages. B cell zone and T cell zone areas captured by high power images on right (indicated by Yellow boxes in the low power image). **(G)**Heatmap showing scaled gene-set expression scores (AddModuleScore) using reference IL10 or IL4 stimulated macrophage gene-sets (*Xue* 2014).**(H)** Dotplot showing NALT MNP expression of *COL6A1*, *COL6A2, CCL18, ITGB2, LILRB4, MMP9.* Size of point indicates fraction of cells in each group expressing gene, colour of point indicates scaled mean expression of gene in each named cell type group.

Examining subset proportions in nasal tissue, we found an increase in monocytes and moDCs in acute COVID-19 (**Fig 3B**). The CD14+ MoDCs were localised to the subepithelial region (**Fig. 3C, S3B**) and expressed ficolin (*FCN1*) (**Fig 3D**), a transmembrane c-type lectin that mediates CXCL8 upregulation upon pathogen encounter ^27^. A small proportion of moDCs also expressed *CXCL8* (**Fig 3D**) and showed reduced enrichment of antigen presentation genes in acute COVID-19, in contrast to nasal cDCs (**Fig. 3E**). Altogether, this is consistent with a functional switch in moDCs in acute COVID-19 away from antigen presentation and toward neutrophil recruitment.

*C1Q*^hi^ macrophage representation in NALT remained stable in health, acute and convalescent COVID-19, but *FOLR2*+ macrophages were significantly increased in convalescent COVID-19 (**Fig 3B**). Spatially, macrophages were predominantly located in T cell zone, including the FOLRB+ subset (**Fig. 3F, S3B**). These *FOLR2*+ macrophages also showed marked temporal changes in their transcriptional programme between acute and convalescent COVID-19, demonstrating enrichment of IL10- and IL4-stimulated macrophage gene signatures ^28^ in convalescent disease (**Fig. 3G, S3C).** EMT pathway genes highly expressed in *FOLR2+* macrophages included *COL6A* (**Fig. 3H**). Collagen VI is an important structural scaffold in many tissues ^29^, and its expression in convalescent NALT macrophages is consistent with a role in structural repair to enable the on-going generation of adaptive immune responses post-infection. Indeed, these *FOLR2*+ macrophages also expressed the tolerogenic receptor *LILRB4*, matrix metalloproteinases, and *CCL18* (**Fig 3H)**, a pro-fibrotic cytokine associated with the differentiation of ‘M2’ like macrophages ^30^. Flow cytometric analysis confirmed increased expression of CD206, a marker of M2 macrophages, in nasal CD14+ cells in convalescent COVID-19 (**Fig S2E)**.

Altogether, our analysis suggests that NALT myeloid cells play differing roles during the progression of SARS CoV2 infection; infiltrating monocytes and moDCs produce self- and neutrophil-recruiting chemokines to set-up a pro-inflammatory positive feedback loop in acute COVID-19 and localise to the periphery of NALT, forming a defensive shield. Meanwhile, NALT macrophages reside predominantly within the T cell zone, and include a *FOLR2*+ subset that is expanded and transcriptionally activated in convalescent COVID-19, adopting a pro-repair, anti-inflammatory transcriptome, promoting tissue integrity and ensuring the infrastructure for the generation of local adaptive immune responses is maintained.

### NALT CD8 tissue-resident memory T cells exhibit clonal expansion and prolonged activation

We next considered T, NK cell and innate-like lymphocytes in isolation and identified 19 cell types (**Fig. 4A, S4A**). We found distinct clusters of naïve and memory CD4 and CD8 T cells, Tregs, Tfh, innate lymphoid cells (ILCs), gamma delta (γδ) T cells, MAIT cells and three subsets of NK cells marked by expression of CD16, CD56 and/or KLRC3 (**Fig. 4A, S4A**). Overall, CD4 T cells were enriched in nasal samples, comprising 80% of CD3+ cells in NALT (**Fig 4B, S4B,** confirmed by flow cytometry, **Fig S4C**). Tissue resident memory (Trm) CD8 T cells, Tfh cells and ILCs were almost exclusively found in the nasal samples, whereas CD16^+^ NK cells, CD8 T effector cytotoxic (CTL), and CD8 T effector memory cells were predominantly found in blood (**Fig. 4B-C**). PD1-expressing Tfh were mostly located within B cell follicles, whilst CD8 Trm predominantly localised to the sub-epithelium, with scattered cells in B cell follicles and interfollicular regions (**Fig. 4D-F, Fig S4M**). In COVID-19, there was an expansion of NALT CD8 T cells compared with healthy controls, particularly naïve, Tcm and Trm cells (**Fig 4C).** The latter is consistent with studies of Trm in murine lymph nodes, demonstrating that whilst the majority of tissue CD8 Trm remain indefinitely resident, a small number of emigrants re-locate to local draining lymph nodes, becoming broadly distributed to enable rapid expansion and defence upon local re-challenge ^31^.

**Figure 4.**
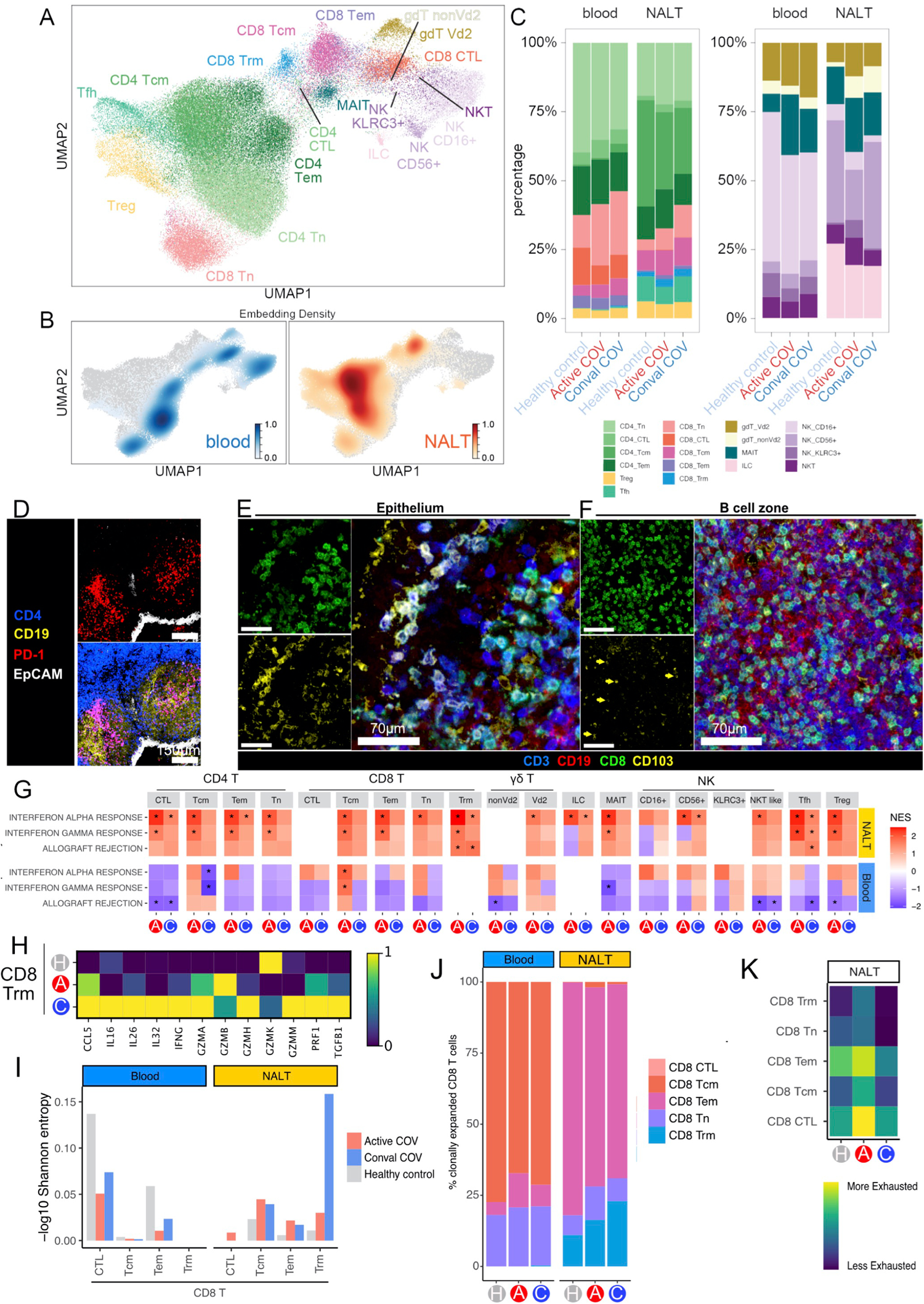
NALT CD8 tissue-resident memory T cells exhibit clonal expansion and prolonged activation. **(A)**UMAP of T cells and innate lymphocytes, subset and re-clustered.**(B)**Embedding density (Gaussian kernel estimation) UMAP of T cells and innate lymphocytes showing scaled density of cells by sample type where 1 = High density of cells **(C)**Stacked bar charts showing proportions of CD4 and CD8 T cells (left paired bar chart) and innate lymphocytes (right paired bar chart) by sample type and disease status. **(D)**Confocal immunofluorescence microscopy of NALT from a convalescent COVID-19 subject showing PD1+ (red) CD4+ (blue) Tfh cells localised to B cell follicles (yellow, CD19). Epithelium shown in white (EPCAM). **(E)**Confocal immunofluorescence microscopy of NALT from a convalescent COVID-19 subject showing CD3+ (blue) CD8+ (green) CD103+ (yellow) Trm cells located in the epithelium/ sub-epithelium. **(F)**Confocal immunofluorescence microscopy image of NALT from a convalescent COVID-19 subject showing CD3+(blue) CD8+ (green) CD103+ (yellow) Trm cells located within B cell follicles (red CD19). Arrows indicate cells that co-express CD8 and CD103.(G)Heatmap of differentially expressed gene enrichment of selected GSEA Hallmark gene sets in T cell and innate lymphocyte subsets by COVID status against healthy control samples. Colour of tile shows normalised enrichment score (NES). Red, Increase relative to healthy controls; Blue, Decrease; * Statistical significance (p-value<0.05 and adjusted p-value <0.1). **(H)**Scaled heatmap showing selected gene transcripts highly expressed in NALT CD8 Trm in Convalescent COVID-19 (C), compared with Healthy control (H) and Active COVID-19 (A) subjects. **(I)**CD8 TCR clonality assessment as diversity (Shannon entropy index) by cell subset and sample type, in Active COVID-19 (red) and Convalescent COVID-19 (blue), and healthy controls (grey). Shannon entropy value has been -log10 transformed so that higher values indicate lower TCR diversity/greater TCR clonality. **(J)** Bar chart showing proportions of clonally expanded CD8 T subsets from single cell TCR analysis, split by sample type. **(K)**Heatmap showing expression (AddModuleScore) of human CD8 T cell viral ‘exhaustion’ signature (*Quigley* 2010) in NALT CD8 T cell subsets.

As with myeloid cells, nasal T/innate lymphocyte subsets demonstrated greater infection-associated transcriptional changes compared to their circulating counterparts, with *‘Interferon alpha response*’ and ‘*Interferon gamma response*’ gene-sets significantly enriched in acute COVID-19 for most T/innate lymphocyte cell types in NALT, with limited/absent enrichment in blood (**Fig. 4G**). In particular, the GC-associated Tfh showed upregulation of interferon-induced anti-viral molecules in acute COVID-19, including *IFITM1* (**Fig. S4F**). In contrast to acute disease, in convalescent COVID-19, ‘*Interferon alpha response*’ gene-set expression returned to healthy control levels in all NALT CD8 T cell subsets except for CD8 Trm (**Fig 4G, S4G**). Indeed, CD8 Trm showed evidence of on-going activation in convalescent subjects with significant enrichment of several immune-relevant genesets (**Fig 4G, S4H**), including cytotoxicity-effector programmes, with *IFNG* and the anti-microbial peptide/cytokine *IL26* among notable leading-edge genes (**Fig. 4H, S4I-J**). IL-26 binds to DNA released from damaged cells to promote damage-associated inflammation ^32^ and may also act directly on epithelial cells, promoting CXCL8 production ^33^. In convalescence, these Trm also expressed tissue repair-associated molecules including *TGFB1* and *IL32*, as well as *IL16* and *CCL5*, T cell recruiting chemokines (**Fig. 4H**). Single cell TCR analysis showed the lowest levels of diversity in the nasal CD8 Trm population in convalescent COVID, as measured by Shannon entropy (**Fig 4I)**, suggestive of clonal expansion. Indeed, comparing healthy control, acute and convalescent COVID-19, a progressive increase in clonally expanded T cells (defined as TCR clonotype with >=2 cells) was uniquely detectable in Trm among nasal CD8 T cells (**Fig 4J)**.

Persistent CD8 T cell activation can lead to exhaustion. Overall, canonical exhaustion-associated genes ^34^ and inhibitory molecules such as *TIGIT*, *LAG3*, *VSIR,* and *PDCD1* (encoding PD1), were upregulated in convalescent compared to acute COVID-19 (**Fig 4K, S4H, K**). Here CD8 Trm again differed from other cytotoxic nasal CD8 T cell subsets, with lower levels of exhaustion geneset enrichment and less expression of *TIGIT and PDCD1* in convalescent samples (**Figs 4K, S4L)**.

Together, our data reveals marked induction of interferon-induced anti-viral programmes in NALT T cells in active COVID-19 infection that rapidly subsides in convalescence. The exception to this was CD8 Trm, which alone showed clonal expansion and evidence of on-going cytotoxic activity in convalescence, as well as production of IFNγ, anti-microbial peptides, and pro-repair molecules. Remarkably, despite this persistent activation, Trm had reduced transcriptional evidence of exhaustion compared with other CD8 T cell subsets. Although Trm have previously been described in human SLOs ^35^, our study provides a unique insight into how NALT Trm respond during viral challenge, suggesting they play parallel roles in defence, the coordination/recruitment of other T cell subsets and in tissue repair.

### Anti-viral responses and GC progression in SARS CoV2 infection

In the B cell compartment, we identified naïve, non-switched and switched memory, FCRL4+ mucosa-associated memory B cells, IgA/IgG plasma cells (PC) and early plasmablasts, as well as distinct clusters of germinal centre (GC)-associated populations, including dark zone, light zone and memory precursors ^4,36,37^ (**Fig. 5A, S5A-B**). Cells annotated as B GC showed expression of canonical GC markers (*AICDA, BCL6*) but failed to enrich for either a light or dark zone signature specifically.

**Figure 5.**
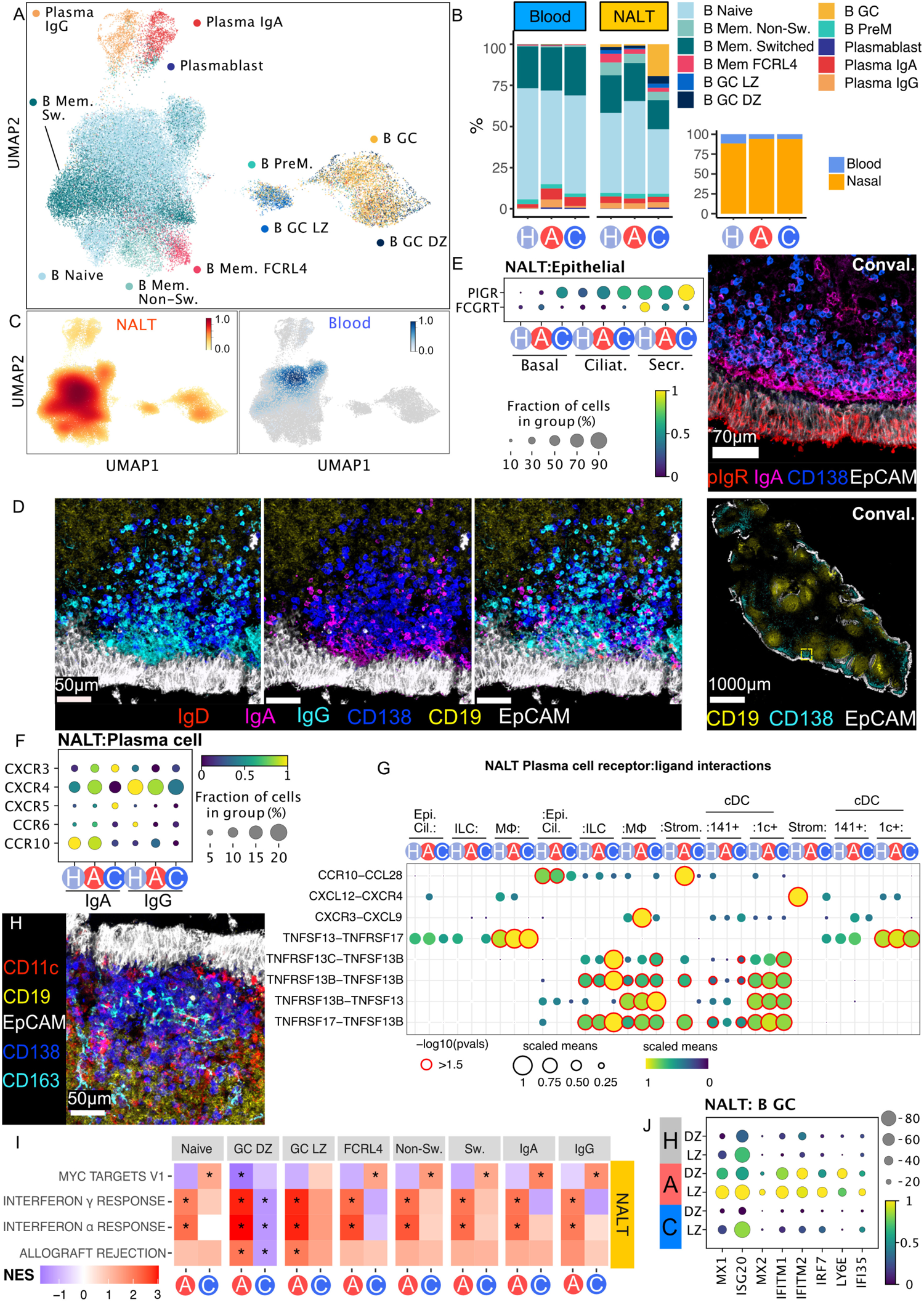
GC and plasma cell expansion in NALT post-COVID infection activation. **(A)**UMAP of B cells and plasma cells, subset and re-clustered. **(B)** Stacked bar chart showing proportions of B and plasma cells in blood and NALT samples by disease status. Nested bar chart (right) shows proportional split of B and plasma cells by disease type and sample type. A, Active COVID-19, C, Convalescent COVID-19, H, Healthy control.**(C)** Embedding density (Gaussian kernel estimation) UMAP of B and plasma cells by sample type where 1 = High density of cells.**(D)**Confocal immunofluorescence microscopy showing subepithelial distribution of CD138+ plasma cells in convalescent COVID-19. Far right – low power image (CD138=cyan; EPCAM=white; CD19= yellow). Left hand higher power panels (CD138=blue; EPCAM=white; CD19= yellow; IgD =red; IgA=magenta; IgG =cyan). **(E)**Expression of NALT epithelial immunoglobulin transporters. Left, dotplot showing PIGR and FCGRT expression in epithelial cell subset populations in NALT by disease state. Size of point indicates fraction of cells in each group expressing gene, colour of point indicates scaled mean expression of gene for each cell type. Ciliat., ciliated epithelial cells; Secr., secretory epithelial cells. Right, Confocal immunofluorescence microscopy of NALT in convalescent COVID-19 subject showing staining of pIgR. CD138, blue; EPCAM, white; IgA, magenta; pIgR, red. **(F)**Dotplot showing selected chemokine receptor expression in plasma cell populations in NALT, split by isotype. Active COVID-19 (red) and Convalescent COVID-19 (blue), and healthy controls (grey). Size of point indicates fraction of cells in each group expressing gene, colour of point indicates scaled mean expression of gene in each named cell type group.**(G)** Dotplot showing CellphoneDB recptor-ligand analysis for NALT plasma cells and ciliated epithelial cells, ILC, macrophages, cDC and stromal cells. Size and colour of dot indicates scaled mean, where the mean value refers to the total mean of the individual partner average expression values in the corresponding interacting pairs of cell types. Red dot border highlight indicates significance (-log10(p-value)>1.5)**.(H)** Confocal immunofluorescence microscopy showing co-localisation of macrophages (CD163, cyan) and conventional dendritic cells (CD11c, red) with plasma cells (CD138, blue) in the subepithelial region of NALT in convalescent COVID-19. EPCAM=white; CD19= yellow. **(I)** Heatmap of differentially expressed gene enrichment of GSEA Hallmark gene-sets in B and plasma cell subsets by COVID status against healthy control samples. Colour of tile shows normalised enrichment score (Red, Increase relative to healthy controls; Blue, Decrease), * p-value<0.05 adjusted p-value <0.1. (J) Dotplot showing expression of selected highly expressed type I interferon response genes in B cell subsets in NALT.

The majority of B cells profiled originated from nasal samples (**Fig. 5B-C**), confirmed by flow cytometry (**Fig. S5C-D)**. The nasal B cell compartment was proportionally expanded in both acute and convalescent COVID-19 compared with healthy controls, with a prominent increase in GC populations in convalescent samples (**Fig 5B)**. FCRL4+ memory cells decreased in COVID-19, and there was also a proportional reduction in nasal switched memory B cells in convalescence compared with acute disease (**Fig. 5B**), confirmed by flow cytometry (**Fig S5D)**.

Spatially, PC were layered along the entire sub-epithelium in convalescence, with many IgG+ and IgA+ PC co-localising here, along with occasional IgD+ PC (**Fig. 5D, S5I**), the latter previously described in tonsils and nasopharynx ^38^. Interestingly, expression of the polyclonal immunoglobulin receptor (PIGR), required for transepithelial shuttling of IgA ^39^, was evident in NALT epithelium and increased in convalescent COVID-19 (**Fig. 5E**), augmenting the capacity to transport IgA generated by adjacent PC into the nasal space. Transepithelial transport of IgG requires the neonatal Fc receptor (FcRn) ^40^. *FCGRT* transcripts were detectable in nasal epithelial cells but did not show an increase post-COVID-19 infection, and indeed, even decreased (**Fig. 5E**), potentially indicating that IgG produced in NALT may bolster systemic rather than nasal IgG. The majority of IgG+ PC, as well as many IgA+ PC, expressed *CXCR4* (**Fig. 5F**), consistent with their co-localisation to a *CXCL12*-expressing niche. The remaining IgA+ PC expressed *CCR10 and CXCR3* (**Fig. 5F**), with NALT stromal cells the only detectable source of *CCL28* (the ligand for CCR10), but myeloid cells the major source of the CXCR3 ligands *CXCL9*, *CXCL10* and *CXCL11* (**Fig. 5G, S5E**). Indeed, macrophages and DCs also expressed *TNFSF13* and *TNSF13B* (encoding APRIL and BAFF) (**Fig. 5G, S5E**), survival factors for plasma cells ^41^, and co-localised with PC clusters (**Fig. 5H, S5J**).

Nasal B and plasmablasts/ cells showed induction of ‘*interferon alpha response’* pathway genes in acute COVID-19 (**Fig. 5I, S5F**). Notably, GC B cells showed substantial expression of several IFN-dependent anti-viral genes, including *MX1, ISG15, IRF7* and *IFITM1/2* **(Fig. 5J, S5G)**. MX1 directly inhibits viral ribonucleoprotein complexes and has been implicated in SARS-CoV2 defence ^42^, and its expression by GC B cells indicates that these specialised adaptive immune cells retain cell-autonomous anti-viral capacity whilst simultaneously differentiating to produce progeny with class-switched, somatically mutated BCRs. Indeed, among immune cells, light zone B cells also showed the highest expression of interferon-induced protein 20 (*ISG20*), an RNA exonuclease endowed with broad antiviral properties ^43^ in acute COVID-19 (**Fig.5J**).

B cell receptor (BCR) analysis showed an increase in CDR3 junctional length in GC B cell populations in convalescent COVID-19 compared with other groups, suggesting increased V(D)J rearrangement and selection in GC structures in convalescent COVID (**Fig S5H)**. Gini index (a measure of clonal selection) was also significantly increased in convalescent COVID-19 (**Fig 6A)**. Analysis of class switching in expanded clones (>2) showed a marked expansion of IgM+ clones (predominantly non-switched memory B cell clones) and IgG1 and IgG2 clones (predominantly GC B cells in active COVID-19, with IgG clones more numerous in convalescence (**Fig. 6B**). Thus, class switching to IgG rather than IgA appears to be the favoured local mucosal response to SARS-CoV-2 infection. In acute COVID, we found an expansion of memory B cell clones, with expanded GC clones more prominent in convalescence (**Fig. 6C**).

**Figure 6.**
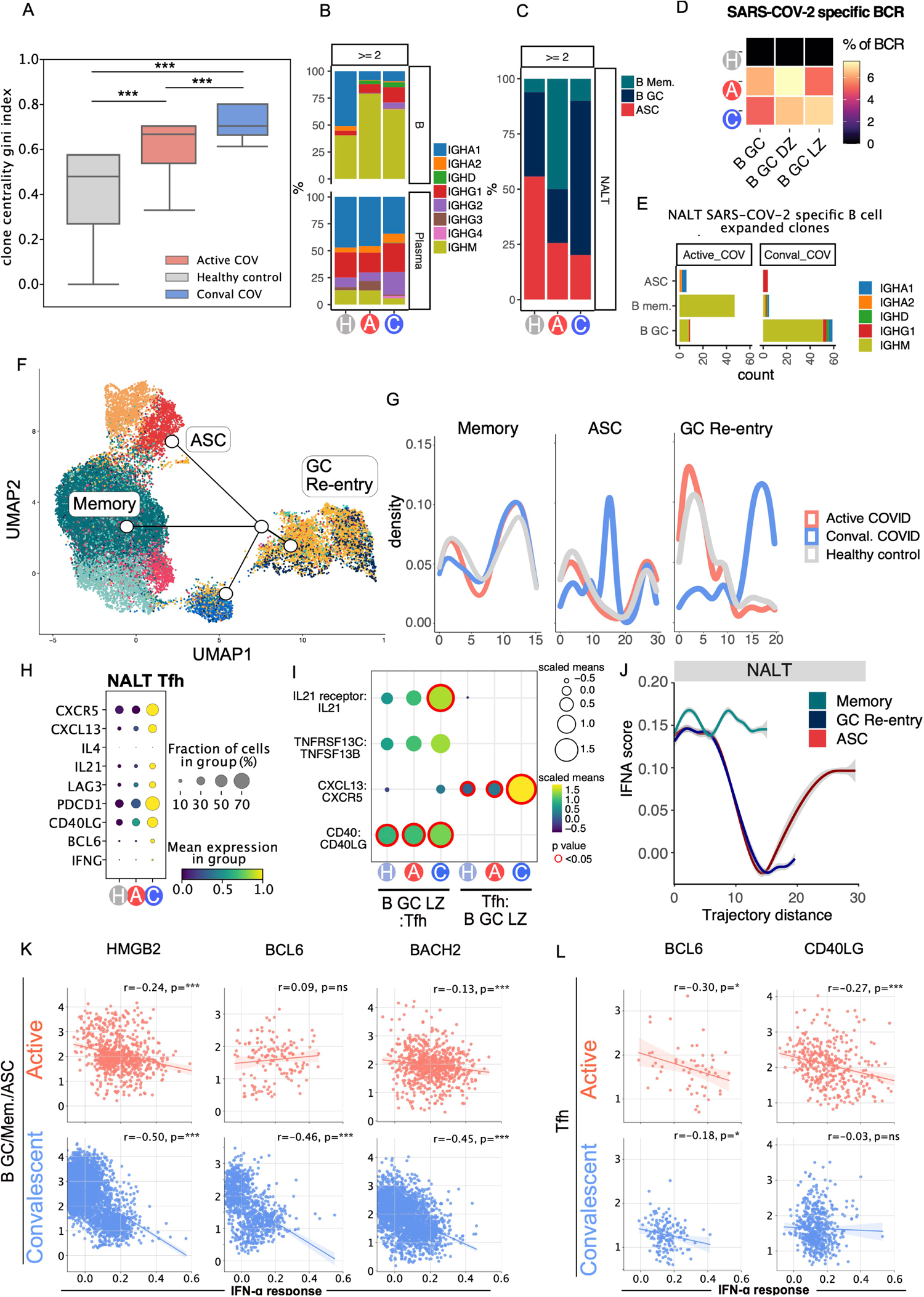
Type I IFN response and GC cell fate and progression. **(A)** Boxplot of NALT BCR Gini centrality index, as a measure of B cell clonality, split by disease state. Gini index values are on a scale from 0-1, where 1 indicates a monoclonal, highly mutated clonal response, ***, p<0.001 (TukeyHSD) **(B)**Bar chart showing proportions of immunoglobulin Heavy chain isotypes in expanded NALT BCR clones (clone size ≥2), by cell type and disease state. Active COVID-19 (pink) and Convalescent COVID-19 (blue), and healthy controls (grey).**(C)**Bar chart showing proportions of memory, GC and antibody secreting cells in expanded NALT BCR clones.**(D)**Heatmap showing SARS-COV-2 specific BCRs as a percentage of total sequenced single cell BCRs in germinal centre B cells, by condition. B GC, Germinal centre B cell; DZ, dark zone; LZ, light zone.**(E)**Stacked bar chart showing counts of expanded SARS-COV-2 specific B cell clones in NALT, by disease state, cell type and isotype. x-axis, cell type; y-axis, count; colour, isotype. ASC, antibody secreting cell; B mem, memory B cell **(F)**B lineage cell slingshot pseudotime trajectories, plotted as UMAP (corresponding to figure 5A). Start point is pinned to B_GZ_LZ (Light zone germinal centre, royal blue). **(G)**Cell density map along pseudotime trajectory distance for memory, GC re-entry and ASC lineages, split by disease state. X-axis, pseudotime trajectory distance, y-axis, density of cells. **(H)**Dotplot showing gene expression in NALT Tfh cells, by disease state, of selected core genes associated with Tfh polarisation and function. Size of point indicates fraction of cells in each group expressing gene, colour of point indicates scaled mean expression of gene in each disease group. **(I)**Dotplot showing CellphoneDB receptor-ligand analysis for NALT B GC LZ and Tfh. Size and colour of dot indicates scaled mean, where the mean value refers to the total mean of the individual partner average expression values in the corresponding interacting pairs of cell types. Red highlight indicates significance (-log10(p-value)>1.5).**(J)** Line graph of GSEA Hallmark interferon alpha response geneset AddModuleScore in memory, GC and ASC B lineage cells along pseudotime trajectory distance for memory, GC re-entry and ASC trajectories (trajectories shown in figure 6D). Memory, Teal; GC Re-entry, Dark blue, ASC, Red. **(K)** Scatterplot with linear regression line plotting expression of GSEA Hallmark interferon alpha response geneset AddModuleScore against HMGB2, BCL6 and BACH2 expression in GC and memory B cells and plasma cells. Linear regression line is shown with a shaded area representing the 95% confidence interval. r, Pearson correlation statistic; p, p-value. *, p<0.05; **, p<0.01; ***, p<0.001; ns, p>0.05. **(L)**Scatterplot with linear regression line plotting GSEA Hallmark interferon alpha response geneset AddModuleScore against *BCL6* and *CD40L* in Tfh.

Next, we sought to identify SARS-CoV-2-specific BCRs, comparing our scBCR data to a reference COVID antibody database (Cov-AbDab) ^44^. SARS-CoV-2-binding BCR sequences were defined as cells with an identical light-chain CDR3 (CDR3-L) sequence and a paired heavy chain CDR3 (CDR3-H) with significant motif similarity to an antibody within the Cov-AbDab (**Fig.S6D**). Putative SARS-CoV-2-binding BCRs (n=768) comprised around 5% of GC B cell BCRs in patients with COVID-19, but were not detectable in control GC B cell BCRs (**Fig. 6D**). Clonal expansion of SARS-CoV-2-specific B cells was also principally evident among GC B cells in convalescent COVID-19 subjects (**Fig. 6E, S6A-C**), supporting the conclusion that this lymphoid tissue generates early local anti-viral antibodies in COVID-19.

The cues that determine which cellular fate a GC B cell adopts and what signals initiate GC exit to a memory B or plasma cell fate in humans remain unclear. Murine studies suggest that this decision is influenced by a combination of BCR antigen affinity, local cytokine environment, and the provision of co-receptor signals by Tfh. A temporal model has been proposed, where the GC response undergoes a switch in its output as it matures, with memory B cells preferred at early timepoints and plasma cells later ^45^. Temporal studies of GC progression in lymphoid organs in humans are lacking, so to investigate this, we used Slingshot pseudotime cell fate trajectory analysis ^46^, anchoring the root of the trajectory in the B GC light zone cluster (B_GC_LZ). This revealed three distinct cell trajectories, one terminating in the antibody-secreting cell cluster (ASC), the second in the switched memory B cell cluster, and third ending in the GC dark zone population, which we termed as a ‘GC re-entry’ fate (**Fig. 6F**). These three trajectories are biologically plausible, and consistent with accepted dogma on GC B cell fate. When comparing disease groups and control, we found a paucity of cells in the terminal stages of the GC re-entry trajectory in acute COVID-19, in contrast to convalescent COVID-19, where many cells were evident in the late GC re-entry trajectory. Cells from convalescent COVID-19 patients also showed an early peak in density in the ASC trajectory compared with acute COVID-19 and control (**Fig. 6G**). Taken together with our analysis of expanded clones, our pseudotime data are consistent with a temporal model of GC progression.

Murine studies suggest that the switch in GC output over time may be mediated by changes in the transcriptional characteristics of Tfh, with the ratio IL21:IL4 decreasing over time, with a switch towards IL4 production. In our dataset, Tfh showed increased expression of *IL21* in convalescent COVID-19 compared to acute disease and control samples, with little *IL4* detectable (**Fig. 6H**). Tfh:Light zone GC B cell interaction prediction using CellphoneDB also showed a temporal increase in several interactions that would promote GC progression, including CXCL13-CXCR5, TNFRSF13C-TNFSF13B, IL21R-IL21 and CD40-CD40LG in convalescent COVID (**Fig. 6I**),

We hypothesised that during viral infection, type I IFNs may provide an environmental cue that influences GC output. Assessment of gene-expression showed persistent induction of type I IFN response genes in cells across the memory B cell trajectory, in contrast to cells progressing along a GC re-entry or ASC trajectory (**Fig. 6J**). Interestingly, in GC B cells, there was a negative correlation between type I IFN response genes and HMGB2 and *BACH2* in SARS-CoV-2 infection, particularly in convalescent disease (**Fig. 6K, S6E-F**). In NALT Tfh, type I IFN response genes showed a negative correlation with *BCL6* in COVID-19 (**Fig. 6L**). Altogether, this is consistent with the conclusion that type 1 interferon may inhibit the progression of the GC reaction, potentially providing an explanation as to why severe COVID-19 is associated with limited GC formation.

## Discussion

Although there is macroscopic involution of the pharyngeal tonsils with age, our study shows that the post-nasal space contains lymphoid tissue in adults that is responsive to viral infection, and represents a feasible method to profile the cellular molecular processes occurring during the generation of adaptive immune responses in living subjects. NALT immune responses are particularly relevant to pathogens that enter via the nasal mucosa, including SARS-CoV-2, and to nasal vaccines ^10,11^. We demonstrate that NALT profiling enables the study of cell types that are central to adaptive immunity, but not present in blood, particularly GC and stromal cells. In addition, the magnitude of transcriptional responses in NALT cells was greater than in circulating cells, emphasising the utility of this method to provide insights into the processes occurring in immune cells during SARS-CoV-2 infection.

Lymphoid tissues contain several myeloid cell populations. We found that monocytes formed a superficial lining in NALT, expressing the antibacterial protein calprotectin ^21^, as well as self- and neutrophil recruiting chemokines. Neutrophils were also evident in this zone, some of which had histone citrullination, required for chromatin decondensation and NET formation ^22^. NETs can also entrap and kill bacteria ^47^, potentially acting together with monocytes to defend the underlying follicles and T cells from nasal bacteria during a period of vulnerability when overlying epithelial cells are damaged or killed by invading virus.

Murine studies show that subcapsular sinus macrophages play important roles in viral defence ^8^. Furthermore, lymphatic sinus-lining macrophages produce IL-18 in response to bacterial infection stimulating innate lymphocytes to produce IFNγ, which in turn enhanced macrophage antibacterial function ^9^. The pharyngeal tonsils do not have afferent lymphatic drainage via a subcapsular sinus, but rather antigens and pathogens come directly from the overlying nasal space. Here we find that human NALT contains a dense network of macrophages, including a *FOLR2*+ population, that expanded in convalescent COVID-19, expressing *IL18,* and pro-repair molecules, including type VI collagen. This is consistent with a role in innate T cell communication and in restoration of lymphoid tissue infrastructure. In acute COVID-19, monocytes and moDCs were recruited to bolster defence, producing self- and neutrophil-recruiting chemokines and localised to form a shield for B cell follicles in the overlying subepithelial region.

Murine models of upper respiratory tract influenza challenge have shown CD8 Trm expansion in the nasal tissue ^48^ and SARS-CoV-2 specific CD8 Trm have also been described in human nasal mucosa ^49^. Our study confirms their presence in NALT and shows that unlike other CD8 T cell subsets, Trm clonally expand and show prolonged cytotoxic gene expression in convalescence, with little evidence of exhaustion. This is in contrast to previous reports suggesting Trm exhaustion in active COVID-19 infection (reviewed in ^50^. Interestingly, NALT Trm also expressed lymphocyte recruiting chemokines and *TGFB1*, suggesting that they play additional roles beyond immediate antiviral defence.

Our longitudinal analysis of GC progression in SARS-CoV-2 infection provides a unique insight into the dynamics of GC responses in humans. Whilst studies in mice suggest a temporal model where memory B cells emerge earlier in responses, this has been difficult to confirm in humans. We found increased representation of memory B cells in expanded clones in acute COVID-19, which decreased in convalescence, and our pseudotime analysis showed a peak in cell density in the ASC trajectory in convalescence. Together, these analyses are supportive of a temporal model of GC progression. We also found that type I IFN may provide an environmental cue that influences GC output, with memory cells showing the highest expression of type I IFN response genes throughout their trajectory.

Here, we show that adults with mild COVID-19 disease do form GCs in response to SARS-CoV-2. This enabled us to probe potential mechanisms of GC collapse described in fatal disease ^19^, and we identify two potential contributing factors; High IFN response gene expression was associated with reduced expression of *BACH2* and *BCL6* in GC B cells and reduced expression of *CD40L* in Tfh. This implies that an excess of type I IFNs, as has been described in severe disease, may inhibit the progression of the GC reaction. In addition, we found evidence of destruction of NALT stromal cells during acute infection, and decreased expression of *CCL19/CCL21*. This is reminiscent of descriptions of a transient loss of CCL19 and CCL21 expression and stromal populations in NALT in murine LCMV models ^51^.

In summary, our study has provided a unique insight into how nasal adaptive immune responses are generated and defended during SARS-CoV-2 infection. Beyond COVID-19, our study provides proof of principle that this sampling strategy will enable long-sought efforts to interrogate longitudinal changes in lymphoid organs in humans in disease and following therapeutic intervention.

## Online Methods

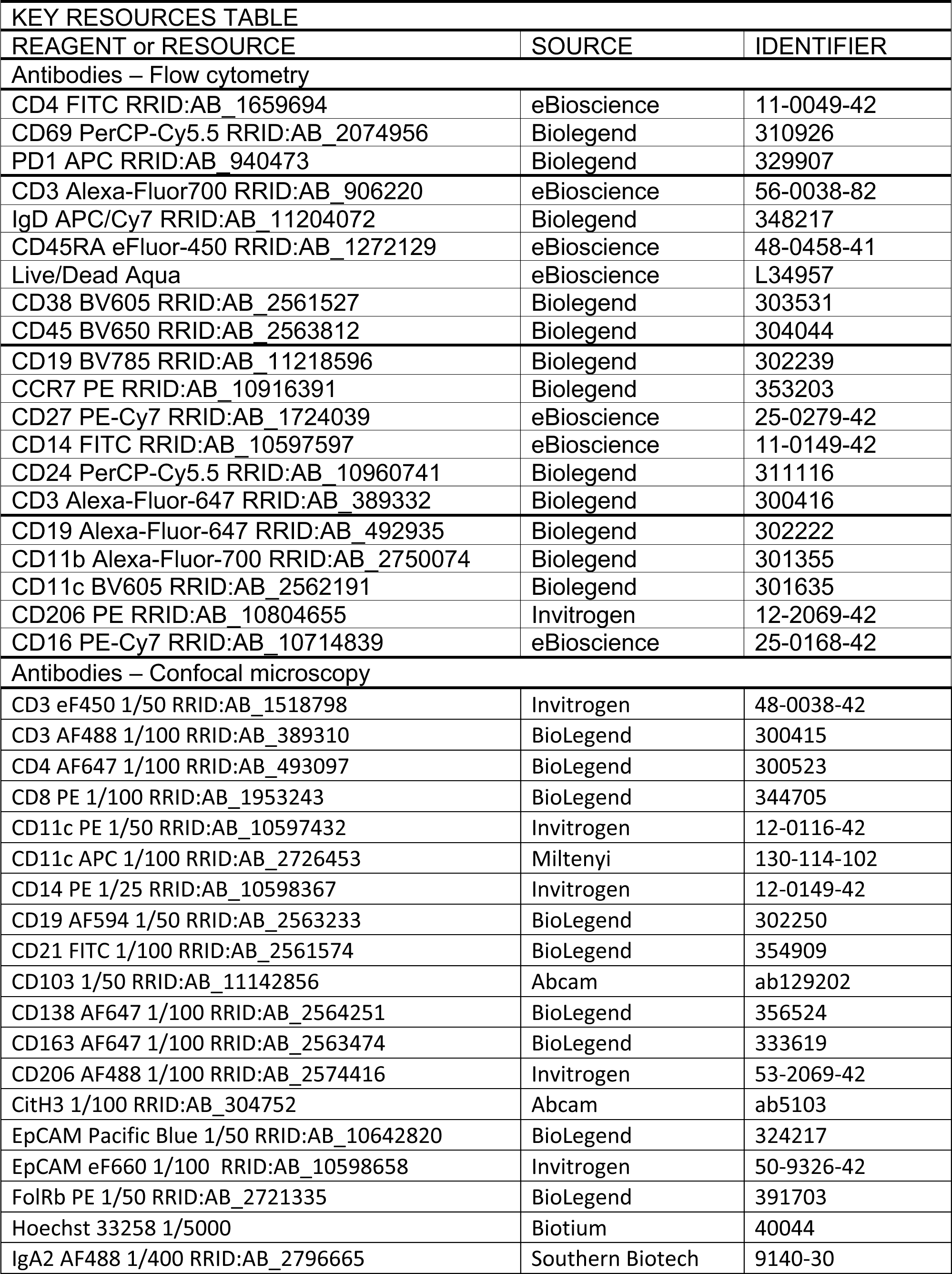
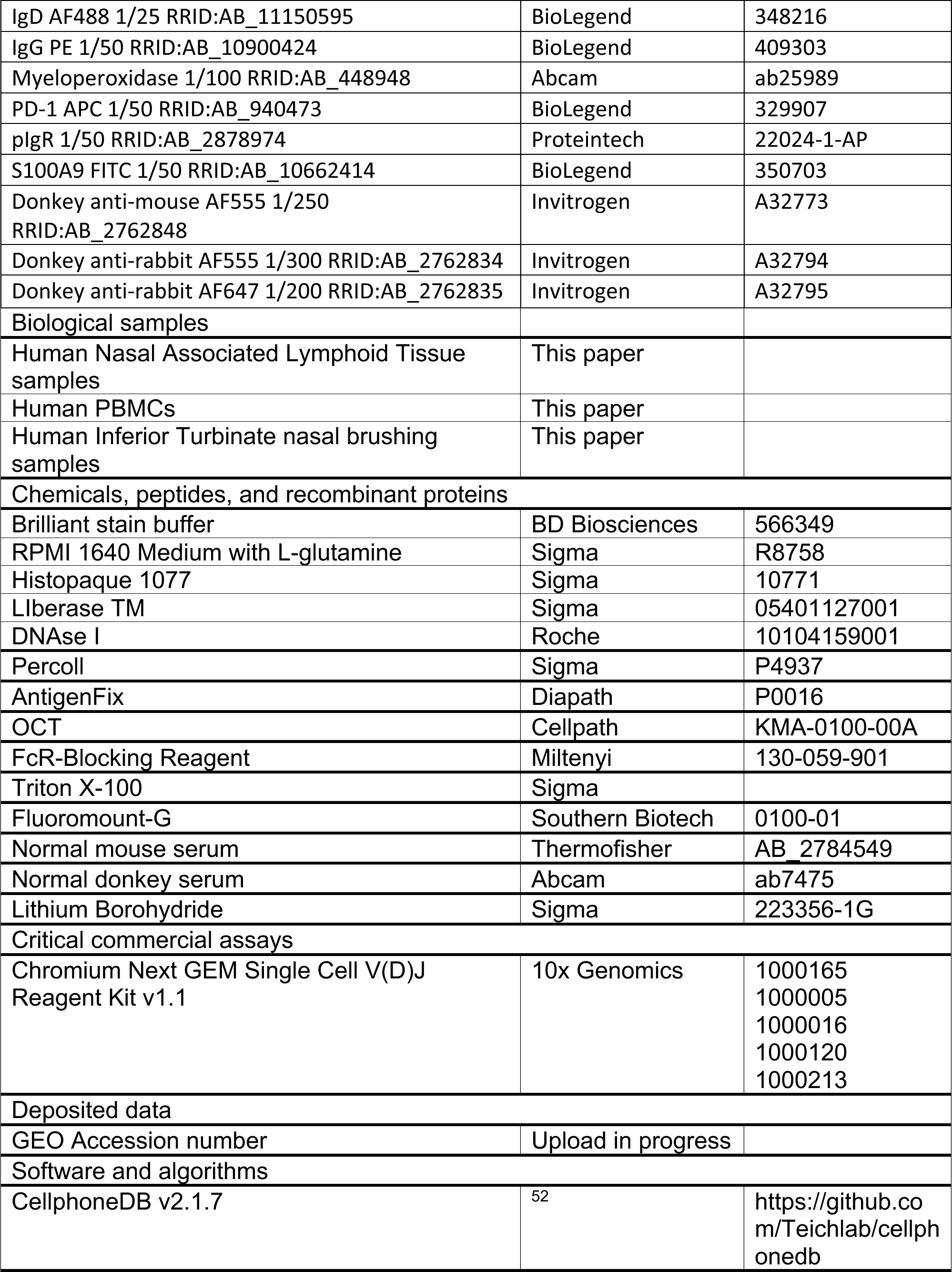
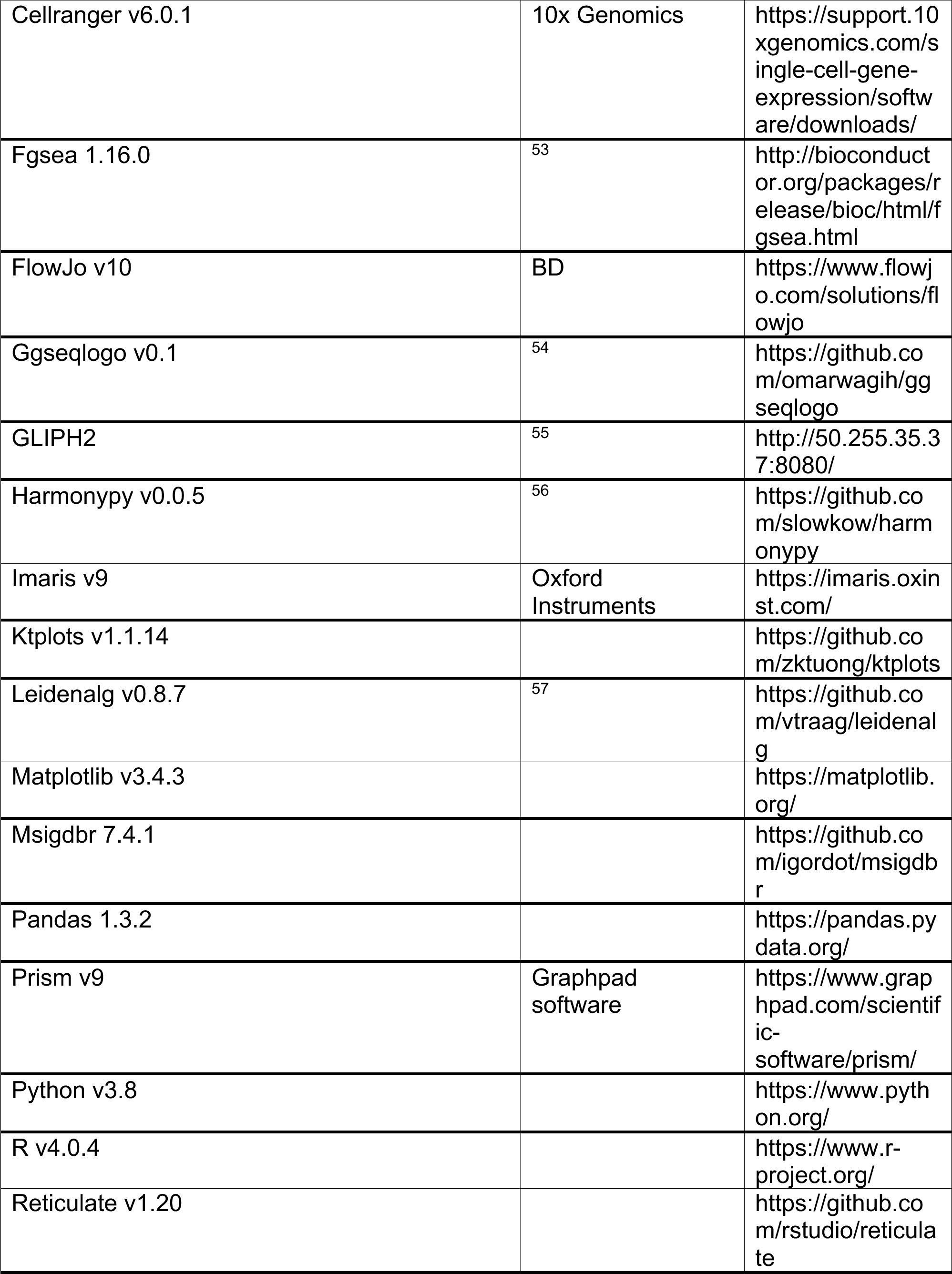
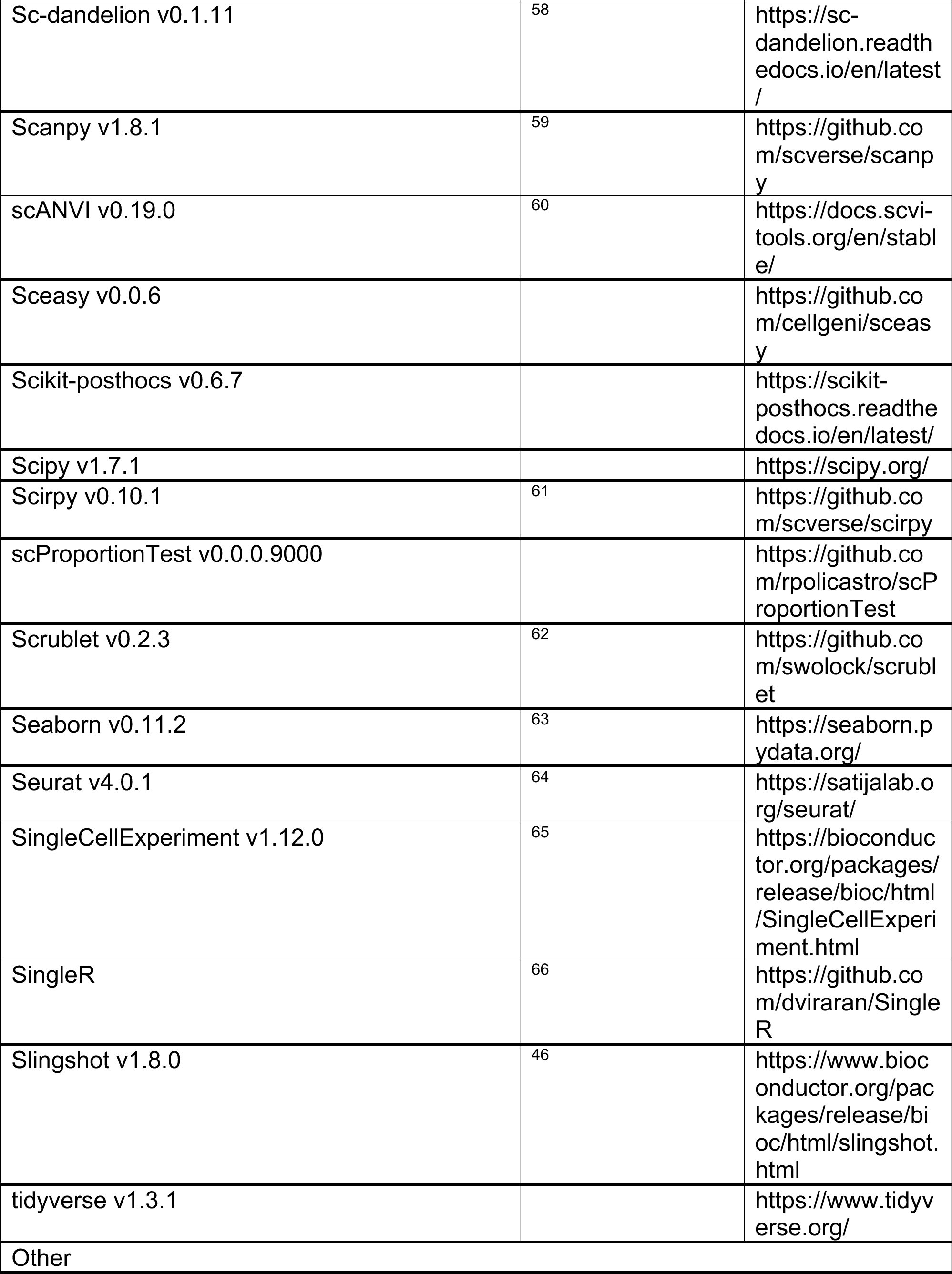
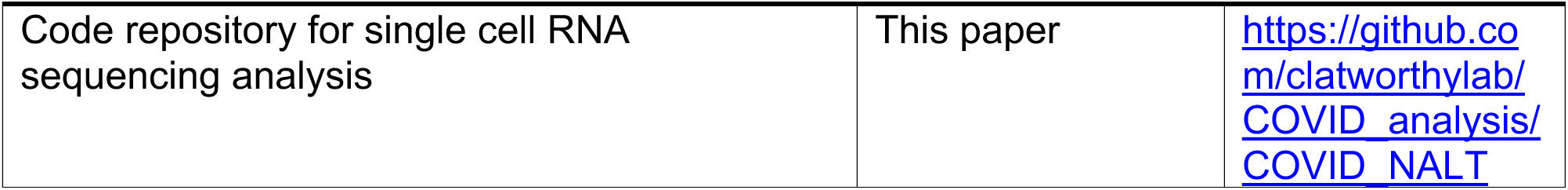

### Human Subject recruitment

COVID-19 subjects were recruited from a single site - Addenbrooke’s hospital, Cambridge, UK, between May 2020 and July 2021. Healthy control subjects were recruited between August 2019 and November 2019 from Cambridge, UK. Ethical approval was given through Cambridge Central Research Ethics Committee (REC reference 08/H0308/267, IRAS project ID: 194217), administered through Cambridge University Hospitals NHS Foundation Trust. All subjects provided informed written consent, and thus patients unable to provide this, including those undergoing invasive ventilation, were excluded. The following subjects were also excluded: <18 years of age, chronic systemic infection (e.g: HIV, Viral Hepatitis), diagnosis of active malignancy, previous head and neck radiotherapy, diagnosis of systemic autoimmune disease (e.g: Rheumatoid arthritis, ANCA-associated vasculitis) or a known nasal/postnasal anomaly precluding access or safe biopsy (e.g: severe septal deviation, nasal vascular malformation). All active COVID-19 subjects, with the exception of subject 1, were recruited within 7 days of their first positive swab (quantitative PCR with reverse transcription – RT-PCR) result and within 14 days of symptom onset. Subject 1 was recruited on the basis of radiological changes consistent with COVID-19, in the presence of positive COVID-19 serology and the onset of symptoms consistent with COVID-19 within the past 14 days. All convalescent COVID-19 subjects were recruited between 21 and 28 days after a positive COVID-19 swab result. No COVID-19 subjects had received COVID-19 vaccine at the time of recruitment.

### Sample collection

Nasal samples were collected by an otolaryngologist under direct endoscopic visualisation. Briefly, the nasal cavity is anaesthetised with topical local anaesthetic and adrenaline (Lidocaine 5%/Phenylephrine 0.5%) administered via spray and neurosurgical patties (Codman). Nasal brushings and micro-curettage were collected from the inferior turbinate under direct vision with the aid of Thudicum nasal specula. The postnasal space (PNS) is inspected trans-nasally with a 0^0^ nasendoscope and 1-3 biopsies taken from the postnasal space mucosa using Tilley-Henkel, Takahashi-fine-cupped or Blakesley forceps, and placed directly into Roswell-Park Memorial Institute 1640 media (RPMI) on ice. Blood samples were collected contemporaneously into Sodium citrate tubes using the Vacutainer system (BD)

### Sample processing to single cell suspension

All samples were processed fresh, within one hour of collection. PNS samples were washed in RPMI, diced and placed into a 15ml Falcon tube with 5ml RPMI and a digestion solution of 62.5μL Liberase^TM^ (Stock conc. 1mg/mL) (Sigma) and 250μL DNAse I (Stock conc. 2.5mg/mL) (Roche). The sample tube was placed into a shaking incubator at 37°C for 30 minutes at 220 RPM. The sample was then pushed through a 70 micron cell strainer (Falcon) into a 50ml Falcon tube, and washed in 50ml RPMI at 500xg for 6 minutes with the brake on. The cell pellet was resuspended in 10ml of 44% Percoll solution and centrifuged at 800xg for 20 minutes at room temperature with no brake. The supernatant was aspirated. If the cell pellet had visible blood staining, 2ml ACK red cell lysis buffer was added for 2 minutes, before quenching with 20ml 1x PBS and centrifuging at 500xg at 4°C for 6 mins (brake on). After aspirating the supernatant, the cell pellet was resuspended in 100μL 1x PBS. Nasal brushing heads were vigorously agitated 15 times in 5ml RMPI collection fluid. The brush head was cut off and left in the collection fluid whilst it was centrifuged at 500xg for 6 minutes at 4°C (brake on). The brush head and supernatant were removed and the cell pellet resuspended in 100μL. Nasal micro-curettes were agitated in 5ml RPMI collection fluid before centrifugation at 500xg for 6 minutes (4°C brake on). The resulting cell pellet was resuspended in a digestion fluid of Liberase/DNAse I at the same concentration as PNS samples, and placed into a shaking incubator for 20 minutes at 37°C, 220 RPM. Following digestion, samples were disassociated by pipette mixing, and washed at 500xg for 6 minutes (brake on) and resuspended in 100μL 1x PBS. 18ml blood from sodium citrate collection tubes was diluted 1:1 with 1x PBS. 15ml Histopaque 1077 (Sigma) was added to each of 2x 50ml Falcon tubes, with the blood-PBS mixture layered on top. Each tube was centrifuged at 800xg for 20 minutes at room temperature (low brake). The resulting PBMC layer was aspirated with a Pasteur pipette, and washed twice in 50ml PBS before being resuspended in 500μL 1x PBS.

The samples were counted with Trypan blue staining to assess cell counts and viability, prior to downstream applications.

### Single cell RNA-sequencing library generation

PNS samples were pooled with curettage and brushing cells at a ratio of 2:1:1. Where nasal curettage and/or nasal brushing cell counts were insufficient or unavailable, only PNS samples were used(Fig S1). Blood and nasal samples were diluted to a concentration of 2×10^6^ cells/ml and loaded onto 10x Chromium controller in parallel lanes with a targeted cell recovery of 14000 cells. Single cell libraries + TCR and BCR VDJ libraries were created using Chromium Next GEM Single cell 5’ kit v1.1 (10x Genomics), in accordance with the manufacturer user guide. Samples were sequenced using a Novaseq 6000 S4 system (Illumina).

### Flow cytometry

After Single cell RNA-seq GEM creation, remaining single cell suspensions were diluted to a concentration of 2×10^6^ cells in 100μL 1x PBS. If this concentration was not achievable, all remaining cells were used. Cells were blocked with 5μL FcR blocking reagent (Miltenyi-Biotech) before undergoing surface staining with fluorophore labelled antibodies as outlined in methods key resources table. Samples were washed and resuspended in 1x PBS and run on Fortessa II flow cytometer (BD). Data was gated in to populations using FlowJo (Treestar/BD). All populations used in this publication gated on a starting population of CD45+ Live, Singlets.

### Immunofluorescence microscopy

Post-nasal space biopsies were collected as described above and fixed for 15-30 minutes (depending on tissue size) in AntigenFix (DiaPath), followed by 8 hours in 30% sucrose in PBS prior to embedding in OCT (CellPath). 15µm sections were mounted on Polysine slides (VWR) and permeabilized and blocked in 0.1M Tris containing 0.1% Triton (Sigma-Aldrich), 1% normal mouse serum, 1% normal donkey serum and 1% BSA (R & D Systems) for 1h at RT. Samples were stained for 2h at RT (primary antibodies) or 1h at RT (secondary antibodies) in a humid chamber with the appropriate antibodies, washed 3 times in PBS and mounted in Fluoromount-G (Southern Biotech). Images were captured using a TCS SP8 (Leica) inverted microscope, on a 40x 1.1NA water immersion objective. Raw imaging data were processed using Imaris (Bitplane/Oxford Instruments).

Iterative staining of post-nasal space biopsies was performed as described by Radtke *et al*.^67,68^. Staining and imaging was performed as described above. After imaging, the coverslip was removed and the slides washed 3 times in PBS to remove any mounting medium. Bleaching of the fluorochromes was achieved using a 1mg/mL solution of lithium borohydride (Acros Organics) in water for 15 minutes at RT. The slides were then washed 3 times in PBS prior to staining as previously described with a different set of antibodies. The process was repeated up to 7 times. Raw imaging data were processed using Imaris (Bitplane/Oxford Instruments) using either Hoechst or CD19 as fiducial for alignment of subsequent images.

### Quantification and Statistical Analysis

#### Single cell RNA-sequencing data processing

Sequenced data was aligned to hu38 human reference genome using Cellranger v6 (10x Genomics). Removal of ambient RNA and assigning of droplets was undertaken using Cellbender ^69^. Automated doublet detection and filtering was performed using Scrublet ^62^ (p value <0.1), as well as manually by gene co-expression. QC and filtering were performed in Scanpy ^59^, using the sc-dandelion pre-processing module (dandelion.pp,recipe_scanpy_qc) with max. genes=6000, min. genes = 200 and a Gaussian Mixture Model distribution to determine mitochondrial filtering values. Batch correction was performed with Harmony ^56^, and Leiden clustering was performed. Cell labels were assigned manually by gene expression profiles (**Fig S1C**), with iterative rounds of sub-clustering to assign cell sub-type labels. Assigned clusters labels were checked against signatures of flow sorted sequenced immune cells ^70^ using SingleR ^66^. Differential gene expression was performed by cell type and condition using FindMarkers in Seurat ^71^, (logfc.threshold=0.1, Min.pct = 0.1, min.cells.group = 10). Gene Set Enrichment Analysis (GSEA) ^72^ was performed on lists of differentially expressed genes using fgsea ^73^ (minSize. = 15) and plotted in ggplot2, with p-values calculated by permutation using fgsea package intrinsic functions. Gene-set expression scores were calculated as the difference between the average expression levels of each gene set and randomly sampled pool of all (control) genes for each cell using scanpy.tl.score_genes: an implementation of Seurat’s AddModuleScore function ^74^. Interferon-alpha response gene expression correlations were calculated using a Pearson correlation coefficient implemented through the scipy package, with two-sided p-values reported. Correlations were calculated on named cell subtypes with named gene expression values >0. Cell proportion plots were performed using ggplot2, with calculation of p-values and confidence intervals using a permutation and bootstrapping approach from the scProportionTest package. Trajectory analysis was performed using Slingshot ^46^. For B cell trajectories, memory B, germinal centre B and plasma cells were subset from the main object and re-clustered. B cell slingshot lineages were inferred with B germinal centre light zone (B_GC_LZ) assigned as the starting cluster. For MNP cell trajectories, monocytes, macrophages and cDCs were subset from the main object and re-clustered. MNP cell slingshot lineages were inferred with classical monocytes assigned as the starting cluster. CellphoneDB analysis was performed as previously described ^52^, with additional plotting of results using ktplots. Single cell TCR analysis was performed using Scirpy ^61^. Single cell BCR analysis was performed using sc-Dandelion ^58^ with VDJ gene reannotation and parsing to AIRR format performed in the sc-Dandelion singularity container (sc-dendelion_latest.sif). Gini index was calculated in sc-Dandelion, with p-values calculated using the Tukey-HSD test implemented through the scikit-posthocs package. For identification of putative BCR sequences with capacity to bind SARS-CoV2 surface antigens, we downloaded CDR3 sequences and germline assignments of known SARS-CoV-2-binding antibodies from the Coronavirus Antibody Database (CoVAbDab, updated 26/07/22) ^44^, and filtered for antibodies of B cell origin. We defined putative SARS-CoV-2-binding BCR sequences in our data set as B cells with complete CDR3 light-chain (CDR3-L) match and significant motif similarity for CDR3 heavy-chain (CDR3-H) compared to the CoVAbDab database. Motif enrichment analysis between BCR CDR3-H sequences in our data and CoVAbDab was performed using GLIPH2 ^55^. First, we compiled a reference database of naïve BCR CDR3 sequences using published CDR3 sequence from Covid-naïve B cells from multiple sources, for the GLIPH2 pipeline. Briefly, 1,437,743 CDR3-H sequences from bulk BCR-sequencing (downloaded from iReceptor gateway 30/07/22) ^75^ and 55,444 CDR3-H sequences from three single-cell GEX + V(D)J-seq studies ^58,76,77^ of control Covid-naïve B cells were included in the reference. For GLIPH2 analysis, we considered only motifs that were over 6 residues in length. We assessed motifs biased towards the target data (CoVAbDab + BCRseq V(D)J data combined) versus the naïve reference data (Fisher exact test p < 0.05), to identify sequences enriched in our cohort. Shared motifs were visualised using ggseqlogo ^54^. We recombined CDR3-H and CDR3-L sequences using their single-cell barcodes and filtered for single cells where both CDR3-H+L sequences matched the corresponding CoVAbDab antibody.

Harmonization and label transfer with a publicly available airway brushing dataset was performed with scANVI^60^, using seed labels from the annotated NALT and PBMC sample cell type labels.

#### Data Availability

All single cell gene expression and VDJ sequencing data has been uploaded to the Gene Expression Omnibus, accession [Upload in progress] and will be freely available on publication.

#### Code Availability

Datails of packages and versions can be found in the online methods key resources table. Code to reproduce the analysis in the manuscript has been deposited in the Clatworthy lab Github repository (https://github.com/clatworthylab/COVID_analysis/COVID_NALT/) and will be freely available on publication.

## Glossary of terms

AMP: Antimicrobial peptide
B GC: Germinal centre B cell
B GC DZ: Dark Zone Germinal Centre B cell
B GC LZ: Light Zone Germinal Centre B cell
B Mem: Memory B cell
B Mem Sw: Switched Memory B cell
cDC: Conventional dendritic cell
CTL: Cytotoxic T cell
gdT/γδT: Gamma-delta T cell
ILC: Innate lymphoid cell
MAIT: Mucosal-associated invariant T cell
MΦ: Macrophage
MNP: Mononuclear phagocyte
MoDC: Monocyte-derived dendritic cell
NES: Normalised Enrichment Score
NK: Natural killer cell
pDC: Plasmacytoid dendritic cell
Tcm: T central memory cell
Teff: T effector cell
Tem: T effector memory cell
Tfh: T follicular helper cell
Tmem: Memory T cell
Tn: Naive T cell
Treg: T regulatory cell
Trm: T resident memory cell
UMAP: Uniform manifold approximation and projection

