## Supplementary for "Temporal profiling of human lymphoid tissues reveals coordinated defence to viral challenge"

**This supplement includes:**

Figs. S1 to S6

Supplement to: Temporal profiling of human lymphoid tissues reveals coordinated defence to viral challenge

### Supplementary 1.

## A

| *Subject* | Cohort | Gender | Age | Ethnicity | COVID severity | Days before biopsy: First positive PCR | Days before biopsy: Onset of symptoms | Days before biopsy: Admission to hospital for COVID | Co-morbidities | Smoking status | COVID therapy | Peak $O_2$ req. | COVID diagnosis | COVID Vaccine | Samples collected: PNS | Samples collected: Brush/MC | Samples collected: PBMC | Notes |
|---|---|---|---|---|---|---|---|---|---|---|---|---|---|---|---|---|---|---|
| *1* | Active COVID | Male | 69 | White-British | Severe | N/A | -11 | -2 | HTN | Ex-smoker | - | 32% | CT | No | ☑ | ☑ | ☑ | |
| *2* | Active COVID | Male | 83 | White-British | Moderate | -3 | -7 | -3 | COPD, IHD, HTN,Asbestosis | Ex-smoker | CS, Baricitinib | 35% | PCR | No | ☑ | ☑ | ☑ | |
| *3* | Active COVID | Female | 29 | White-British | Mild/Asymp. | -1 | N/A | N/A | None | Non-smoker | - | N/A | PCR | No | ☑ | | ☑ | |
| *4* | Active COVID | Female | 90 | White-British | Severe | -2 | -14 | -2 | HTN | Ex-smoker | CS, Remdesivir | 60% | PCR | No | ☑ | ☑ | ☑ | |
| *5* | Active COVID | Male | 76 | White-British | Severe | -1 | -7 | -1 | HTN, DM, PVD | Ex-smoker | - | 40% | PCR | No | ☑ | ☑ | ☑ | |
| *6* | Active COVID | Male | 22 | White-British | Mild | -5 | -6 | N/A | None | Non-smoker | - | N/A | PCR | No | ☑ | | ☑ | |
| *7* | Active COVID | Female | 43 | Asian-other | Moderate | -6 | -8 | 0 | None | Non-smoker | CS | N/A | PCR | No | ☑ | | ☑ | |
| *8* | Active COVID | Male | 24 | Asian-Chinese | Severe | -12 | -13 | -5 | DM | Non-smoker | CS, Tocilizumab | 40% | PCR | No | ☑ | | | |
| *9* | Conval. COVID | Female | 19 | White-other | Mild | -22 | -19 | N/A | None | Non-smoker | None | N/A | PCR | No | ☑ | ☑ | ☑ | |
| *10* | Conval. COVID | Female | 23 | White-British | Mild | -21 | -20 | N/A | None | Non-smoker | None | N/A | PCR | No | ☑ | | ☑ | |
| *11* | Conval. COVID | Female | 24 | White-British | Mild | -28 | -28 | N/A | None | Non-smoker | None | N/A | PCR | No | ☑ | | ☑ | |
| *12* | Conval. COVID | Male | 20 | Asian-British | Mild/Asymp. | -26 | N/A | N/A | None | Non-smoker | None | N/A | PCR | No | | | ☑ | |
| *13* | Conval. COVID | Male | 19 | White-British | Mild | -28 | -28 | N/A | None | Non-smoker | None | N/A | PCR | No | ☑ | | ☑ | |
| *14* | Healthy control | Male | 23 | White-British | N/A | N/A | N/A | N/A | None | Non-smoker | N/A | N/A | N/A | No | ☑ | | ☑ | Sample pre-Dec. 2019 |
| *15* | Healthy control | Male | 26 | White-British | N/A | N/A | N/A | N/A | None | Non-smoker | N/A | N/A | N/A | No | ☑ | | ☑ | Sample pre-Dec 2019 |
| *16* | Healthy control | Male | 35 | White-British | N/A | N/A | N/A | N/A | None | Non-smoker | N/A | N/A | N/A | No | ☑ | | | Sample pre-Dec 2019 |
| *17* | Healthy control | Female | 76 | White-British | N/A | N/A | N/A | N/A | None | Non-smoker | N/A | N/A | N/A | No | ☑ | | ☑ | Sample pre-Dec 2019 |
| *18* | Healthy control | Male | 91 | White-British | N/A | N/A | N/A | N/A | PVD, AF | Smoker | N/A | N/A | N/A | No | ☑ | | ☑ | Sample pre-Dec 2019 |
| *19* | Healthy control | Male | 85 | White-British | N/A | N/A | N/A | N/A | None | Ex-smoker | N/A | N/A | N/A | No | ☑ | | ☑ | Sample pre-Dec 2019 |
| *20* | Healthy control | Male | 30 | Asian-other | N/A | N/A | N/A | N/A | None | Non-smoker | N/A | N/A | N/A | Yes | | ☑ | | No previous symptomatic COVID-19 infection |
| *21* | Healthy Control | Female | 23 | White-other | N/A | N/A | N/A | N/A | None | Non-smoker | N/A | N/A | N/A | Yes | | ☑ | | Lateral flow negative |
| *22* | Healthy Control | Female | 32 | White-other | N/A | N/A | N/A | N/A | None | Non-smoker | N/A | N/A | N/A | Yes | | ☑ | | Lateral flow negative |
| *23* | Healthy Control | Female | 58 | Black-African | N/A | N/A | N/A | N/A | None | Non-smoker | N/A | N/A | N/A | Yes | | ☑ | | Lateral flow negative |

## B

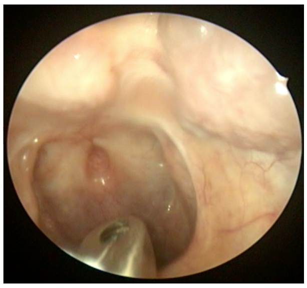

## C

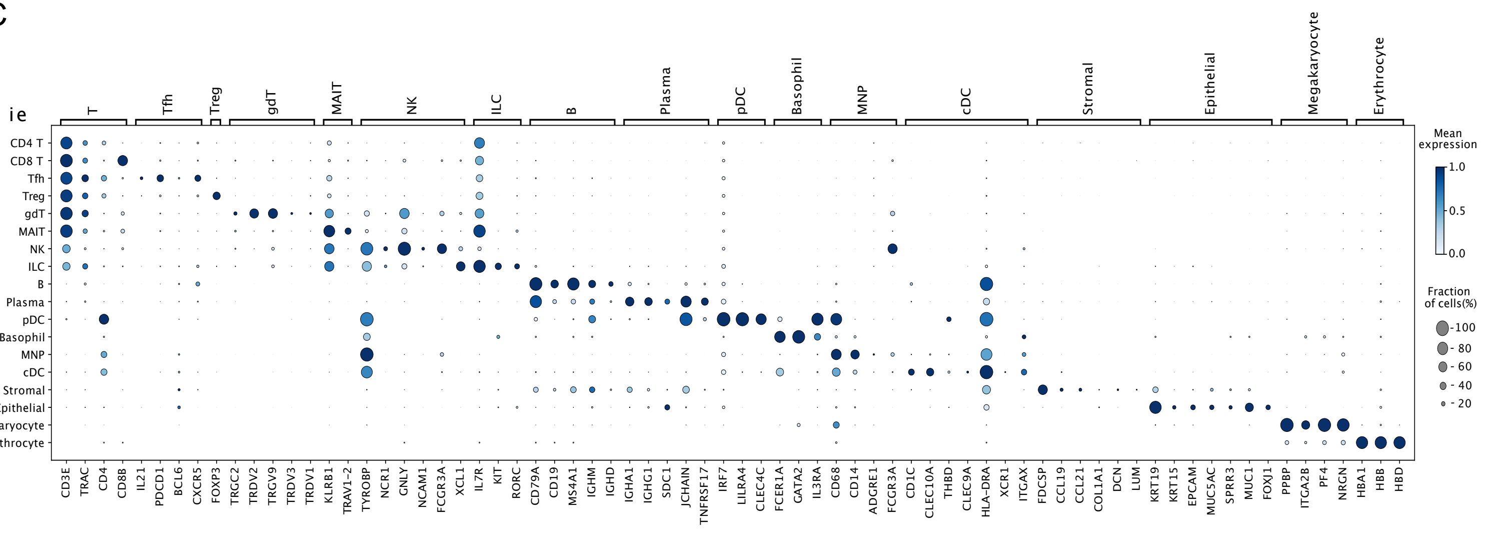

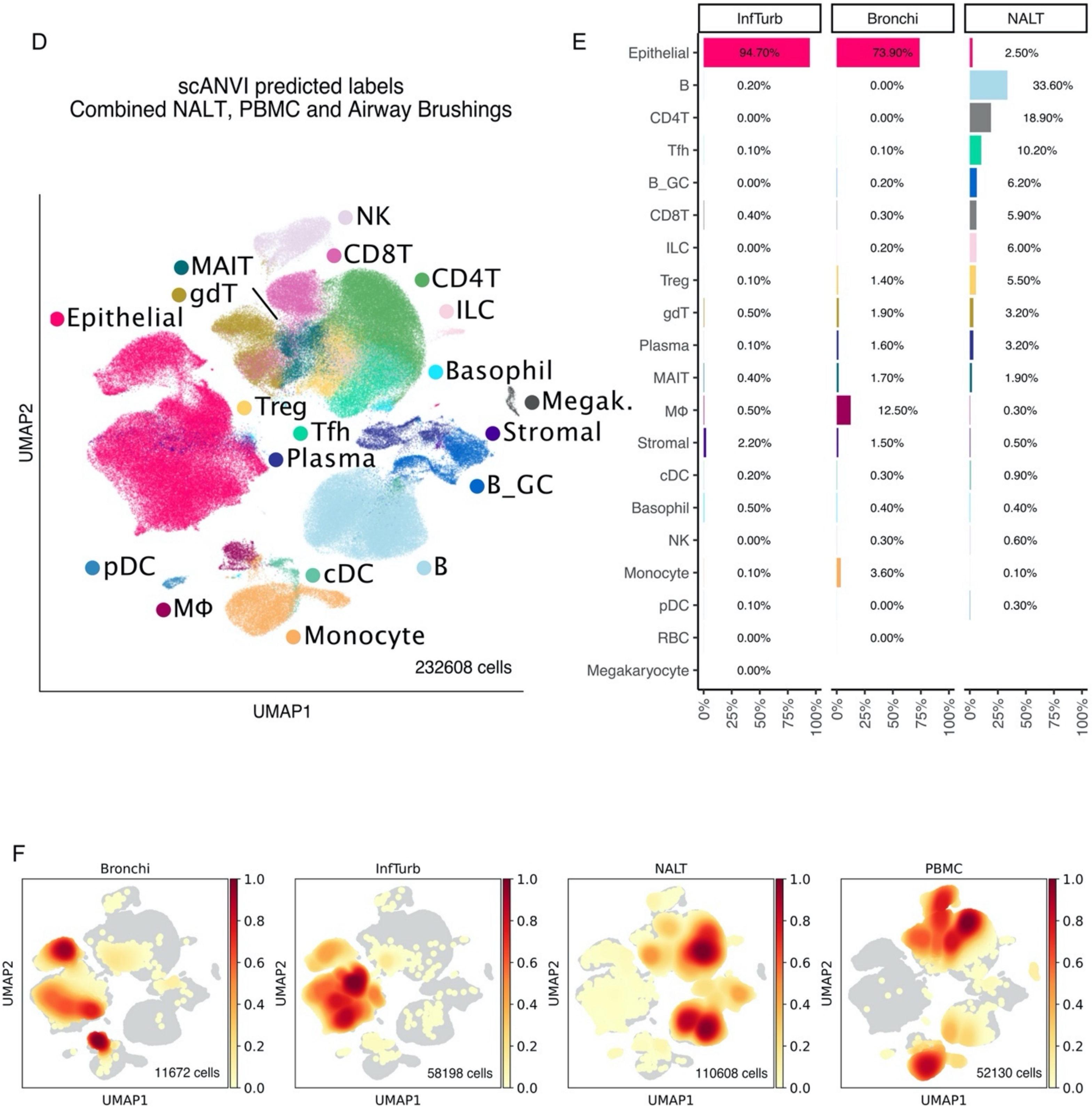
D
scANVI predicted labels
Combined NALT, PBMC and Airway Brushings
NK
CD8T
MAIT
gdT
CD4T
ILC
Epithelial
Basophil
Megak.
Treg
Tfh
Stromal
Plasma
B_GC
pDC
cDC
B
MΦ
Monocyte
232608 cells
UMAP2
UMAP1
E
InfTurb
Bronchi
NALT
Epithelial
B
CD4T
Tfh
B_GC
CD8T
ILC
Treg
gdT
Plasma
MAIT
MΦ
Stromal
cDC
Basophil
NK
Monocyte
pDC
RBC
Megakaryocyte
94.70%
73.90%
2.50%
33.60%
18.90%
10.20%
6.20%
5.90%
6.00%
5.50%
3.20%
3.20%
1.90%
12.50%
0%
25%
50%
75%
100%
F
Bronchi
InfTurb
NALT
PBMC
11672 cells
58198 cells
110608 cells
52130 cells
UMAP1
UMAP2

**Figure S1:**

**(A)** Table showing characteristics of included subjects. COVID severity scoring by WHO criteria [1]. AF, Atrial Fibrillation; Brush, COPD, Chronic Obstructive Pulmonary Disease; CS, Corticosteroids; CT, Computed Tomography scan; DM, Diabetes Mellitus; HTN, Hypertension; Nasal brushing of inferior turbinate; PBMC, Peripheral blood mononuclear cells; MC, Micro-curettage (of inferior nasal turbinate); PCR, SARS-CoV-2 RT-PCR nasopharyngeal swab test; PNS, Postnasal space; PVD, Peripheral Vascular Disease; N/A, Not applicable.

**(B)** Endoscopic nasal view during an office-based postnasal space biopsy procedure in an awake human subject under topical local anaesthesia. Tilley-Henkel forceps are seen towards the bottom of the image

**(C)** Expression of canonical markers by assigned cell type label. Vertical (Y) axis shows assigned broad cell labels, with genes grouped by canonical expression groups on horizontal axes. Dot size shows percentage of gene expression by cells in group, with colour indicating normalised mean gene expression by cells in group.

**(D)** Uniform manifold approximation and projection of scANVI integration of data and cell type labels from this paper with adult inferior turbinate and bronchial brushing data in SARS-CoV-2 subjects and controls previously published[2]

**(E)** Proportional representation of airway sample data from (D) presented by sampling location and predicted cell type following scANVI integration. Cell type percentages are of total cells by sample location

**(F)** Scanpy embedding density UMAP, showing density of cells by sampling location on UMAP projection in (D). Cell numbers by sample location are printed in the bottom right of the projection.

Supplementary 2

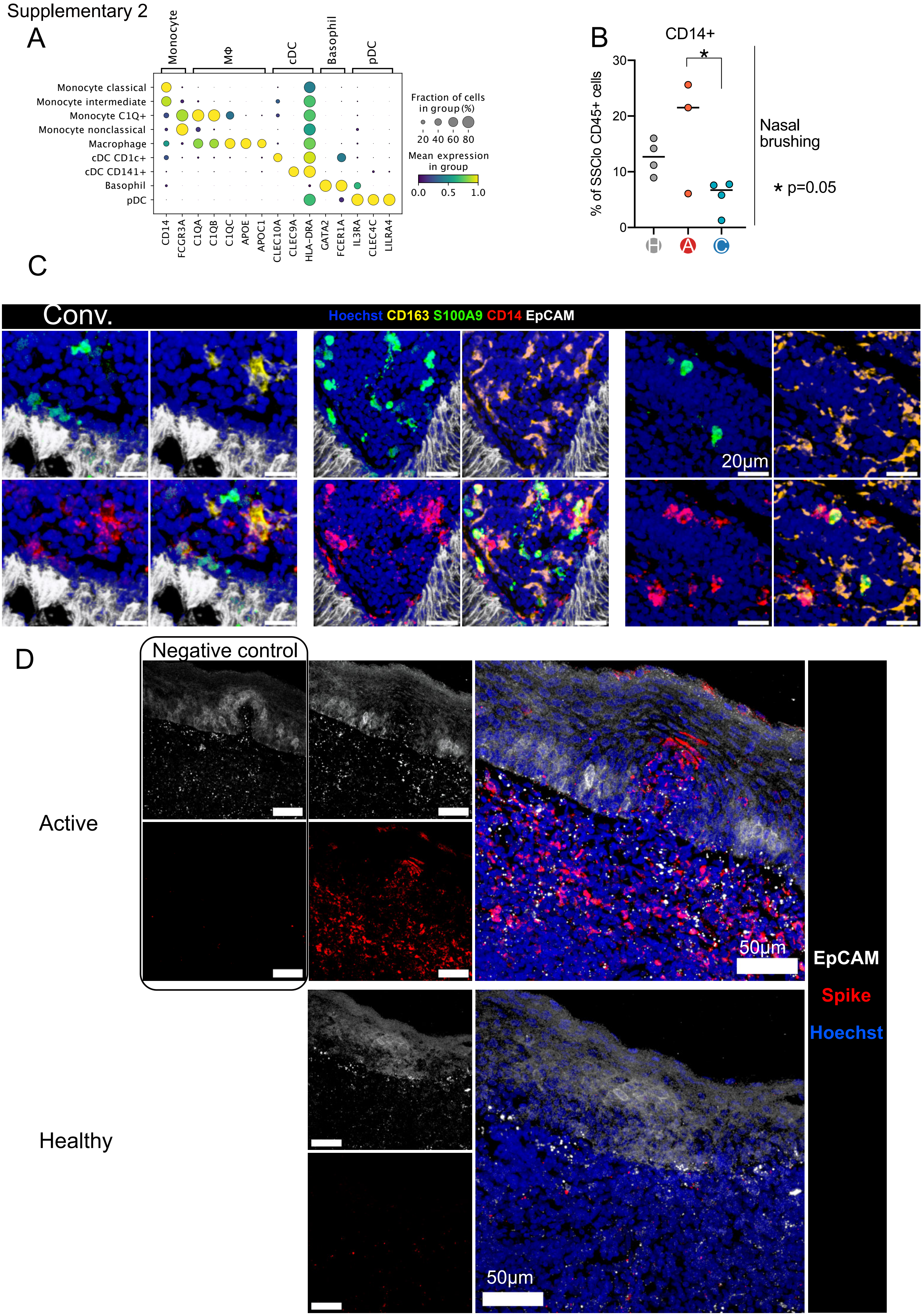

E

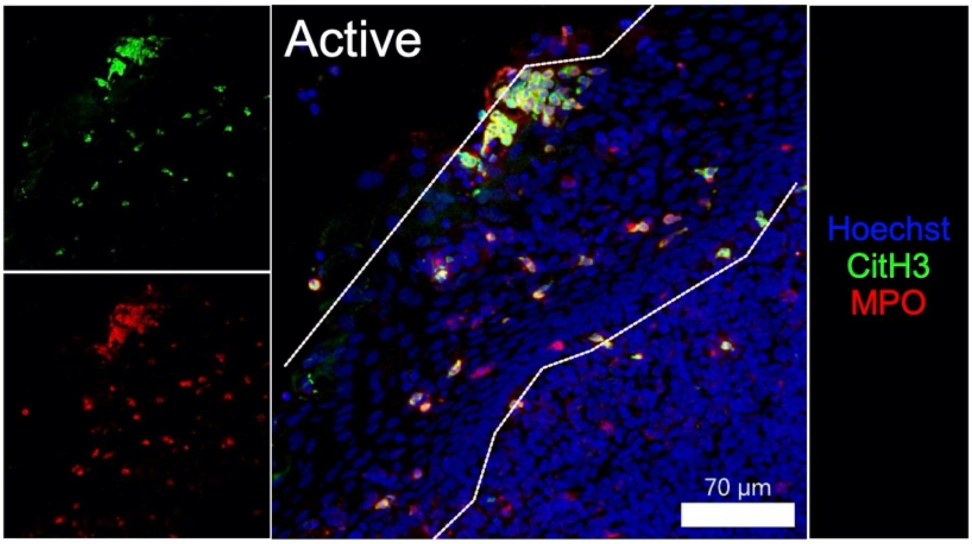
Active
Hoechst
CitH3
MPO
70 µm

F

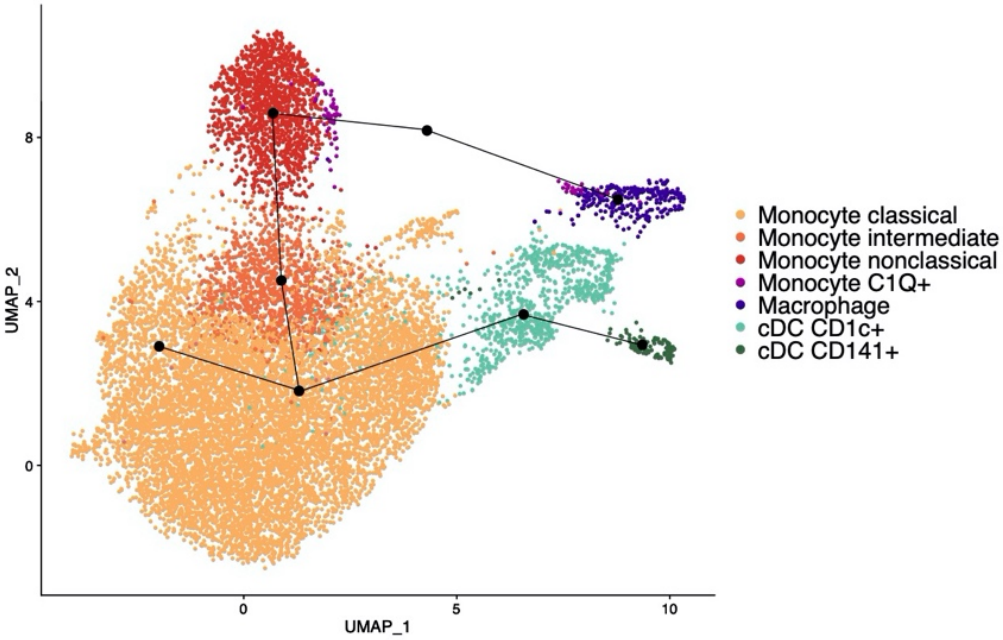
UMAP_2
UMAP_1
8
4
0
0
5
10
Monocyte classical
Monocyte intermediate
Monocyte nonclassical
Monocyte C1Q+
Macrophage
cDC CD1c+
cDC CD141+

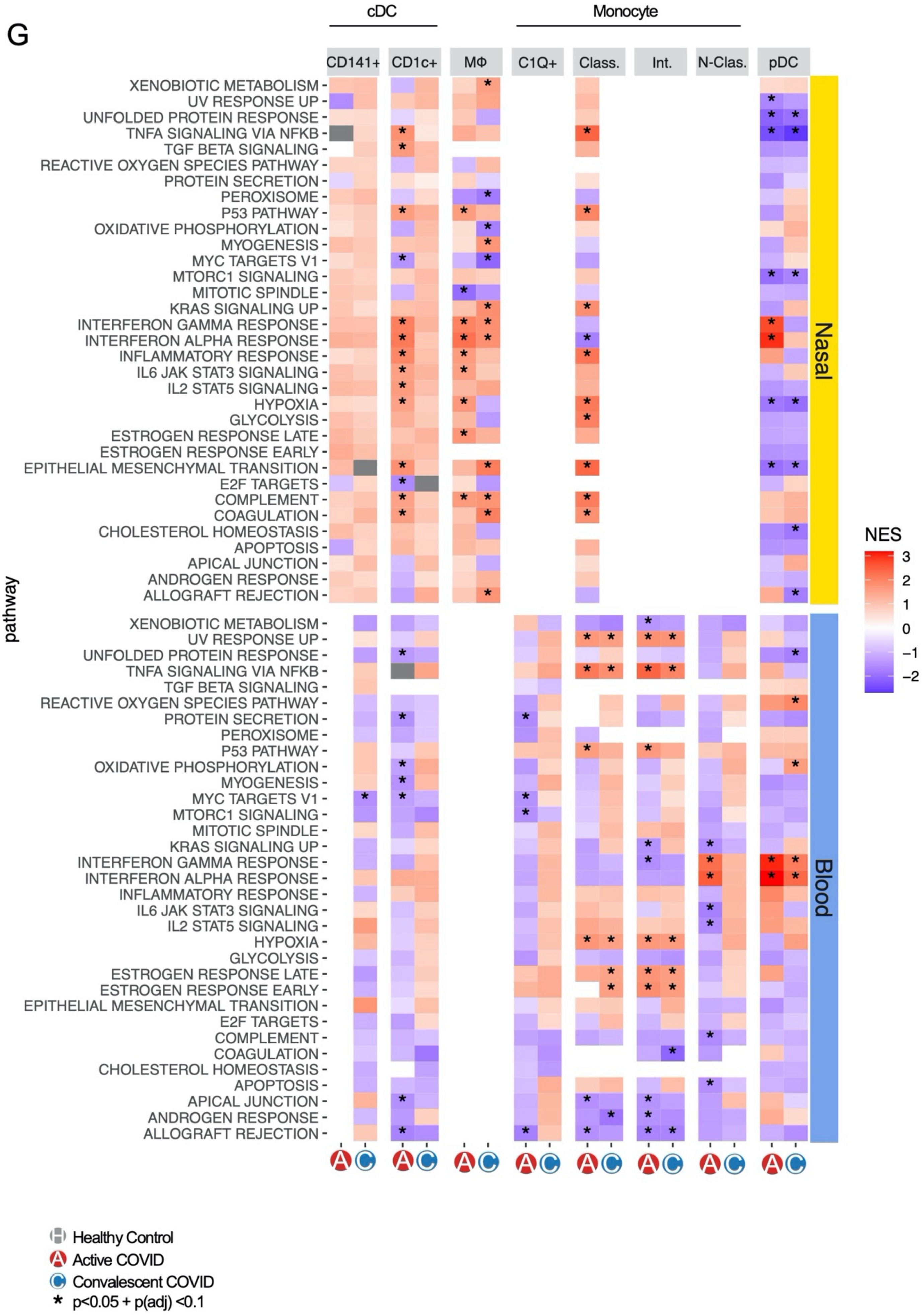
G
cDC
Monocyte
CD141+
CD1c+
MΦ
C1Q+
Class.
Int.
N-Clas.
pDC
XENOBIOTIC METABOLISM
UV RESPONSE UP
UNFOLDED PROTEIN RESPONSE
TNFA SIGNALING VIA NFKB
TGF BETA SIGNALING
REACTIVE OXYGEN SPECIES PATHWAY
PROTEIN SECRETION
PEROXISOME
P53 PATHWAY
OXIDATIVE PHOSPHORYLATION
MYOGENESIS
MYC TARGETS V1
MTORC1 SIGNALING
MITOTIC SPINDLE
KRAS SIGNALING UP
INTERFERON GAMMA RESPONSE
INTERFERON ALPHA RESPONSE
INFLAMMATORY RESPONSE
IL6 JAK STAT3 SIGNALING
IL2 STAT5 SIGNALING
HYPOXIA
GLYCOLYSIS
ESTROGEN RESPONSE LATE
ESTROGEN RESPONSE EARLY
EPITHELIAL MESENCHYMAL TRANSITION
E2F TARGETS
COMPLEMENT
COAGULATION
CHOLESTEROL HOMEOSTASIS
APOPTOSIS
APICAL JUNCTION
ANDROGEN RESPONSE
ALLOGRAFT REJECTION
pathway
Nasal
Blood
NES
3
2
1
0
−1
−2
Healthy Control
Active COVID
Convalescent COVID
* p<0.05 + p(adj) <0.1

**Figure S2**

(A) Expression of canonical gene expression markers by assigned MNP cell label for MNP cell subset. Assigned cell labels on y-axis, x-axis shows canonical marker genes grouped by cell type to which they correspond. Size of point indicates fraction of cells in each group expressing corresponding gene, colour of point indicates scaled mean expression of corresponding gene in each named cell type group.

(B) Flow cytometry of inferior turbinate nasal brushing showing CD14+ cells as a percentage of SSClo CD45+ live single cells. Each point represents a subject. Horizontal lines indicate median value. * p = 0.05 (Tukey HSD test)

(C) Extended images from figure 2E, showing co-expression of CD14 and S100A9 in NALT tissue by immunofluorescence microscopy in a Convalescent COVID-19 subject. Blue, Hoechst, Yellow, CD163, Green, S100A9, Red, CD14, White, EpCAM

(D) Extended images from figure 2F. Immunofluorescence microscopy showing staining of SARS-COV-2 spike protein in NALT epithelium, with negative control (box), from an active COVID-19 subject at high magnification (top). A high magnification healthy control NALT tissue section containing respiratory epithelium is shown to demonstrate a lack of SARS-COV-2 spike protein staining (bottom right). SARS-COV-2 spike protein, Red; EpCAM, White; Hoechst, Blue.

(E) A section of NALT epithelium from a subject with Active COVID-19 infection, imaged by confocal immunofluorescence microscopy, showing neutrophils forming Neutrophil-Extracellular-Traps (NETs). Hoechst, Blue; CitH3, Green; MPO, Red. Respiratory epithelium is indicated by dashed white line.

(F) Slingshot pseudotime trajectories for MNP subset cell types, plotted in UMAP space. Black lines indicate Slingshot lineage trajectories. Trajectory start set to Classical monocyte cluster.

(G) Extended heatmap from figure 2G, showing all significantly enriched Hallmark pathways in MNP cell types following Gene set enrichment analysis of differentially expressed genes between COVID-19 disease groups and healthy control subjects. Only pathways containing at least one significant enrichment ($p<0.05$ and Benjamini-Hochberg adjusted p value $<0.1$) in MNP cell types are shown. Colour indicates normalised enrichment score (NES), with red indicating greater pathway enrichment in disease group compared with healthy controls, and blue indicating increased pathway enrichment in healthy controls than disease group.

#### Supplementary 3

A

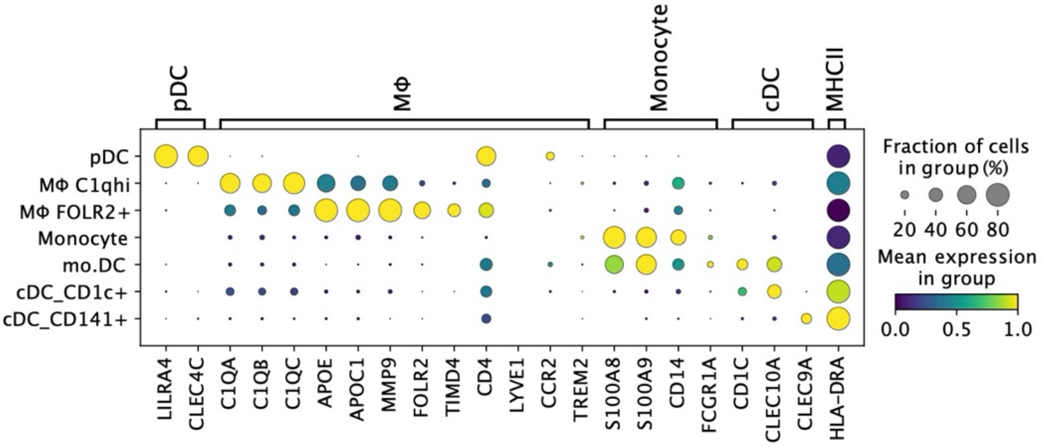


B

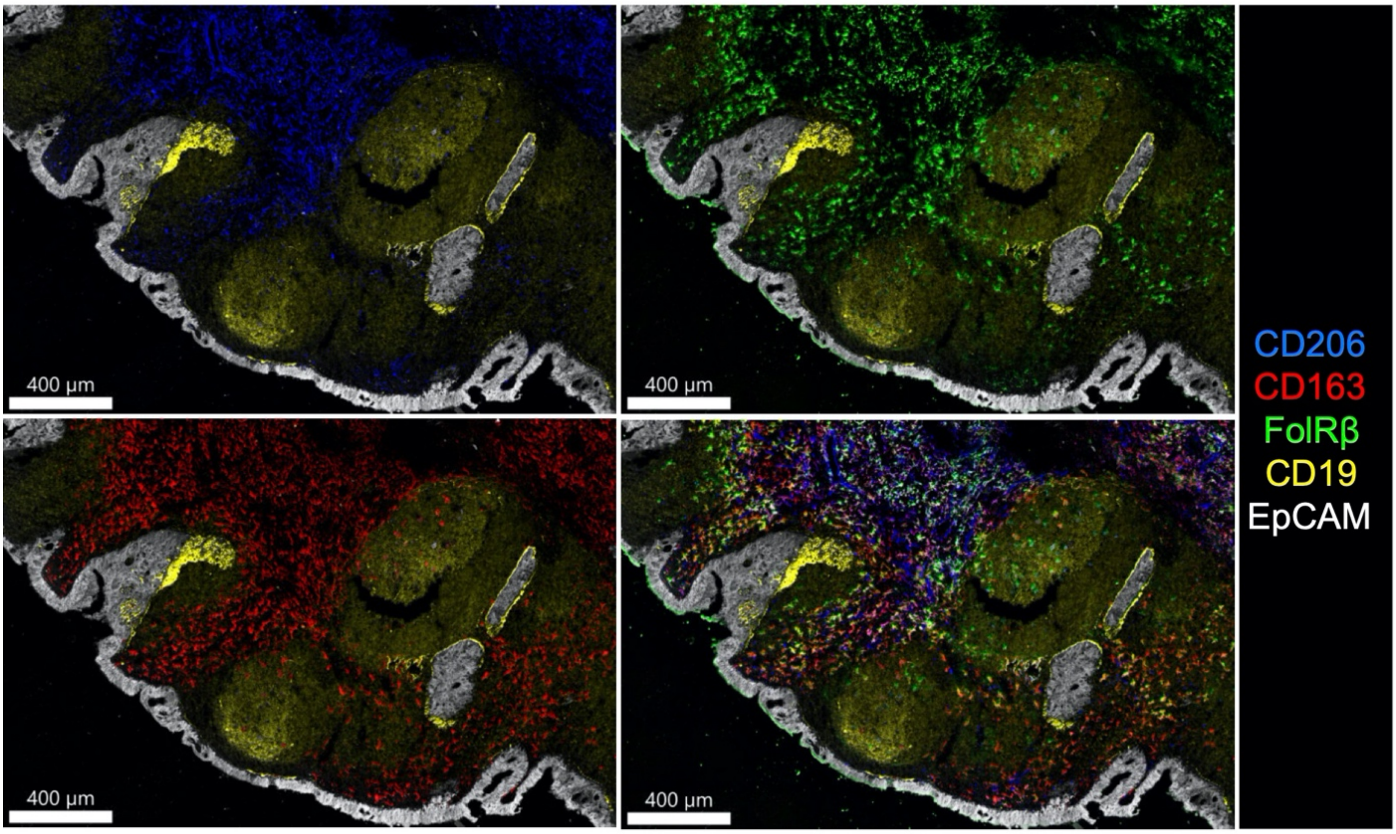

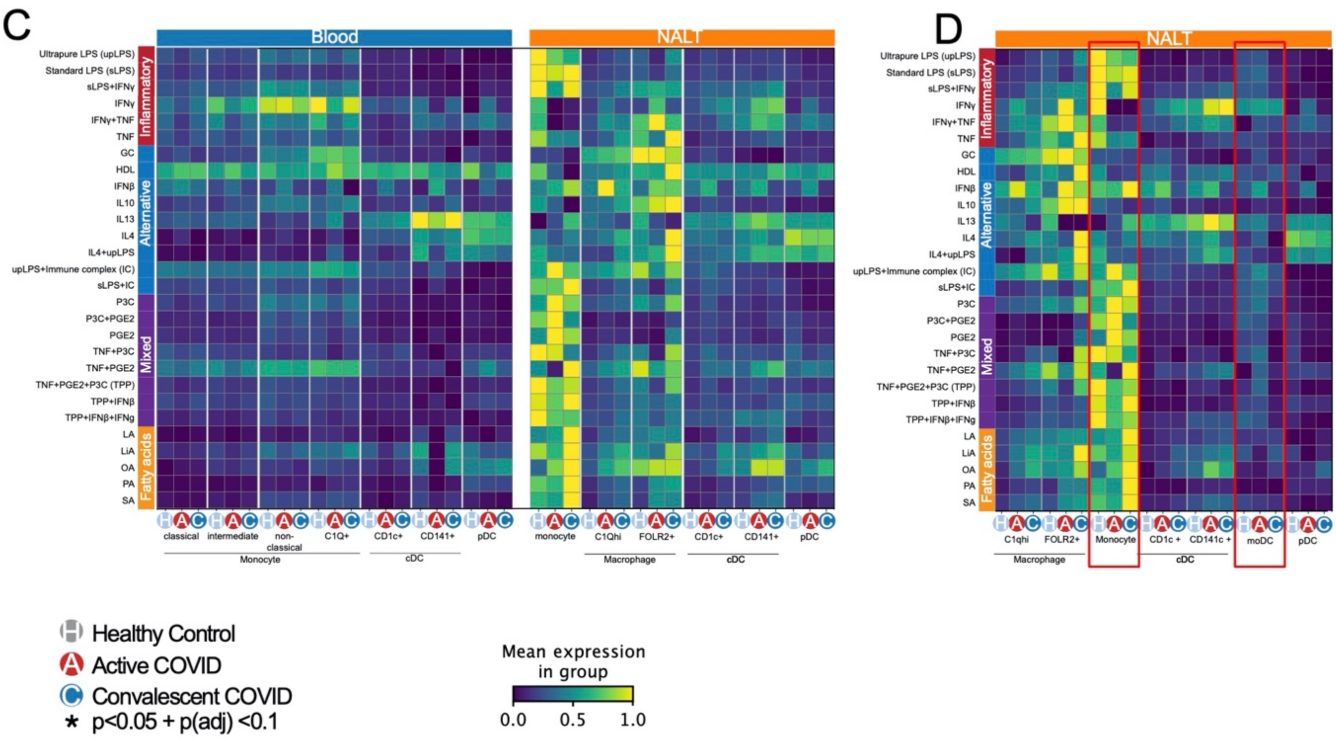
C
Blood
NALT
D
Inflammatory
Alternative
Mixed
Fatty acids
Ultrapure LPS (upLPS)
Standard LPS (sLPS)
sLPS+IFNγ
IFNγ
IFNγ+TNF
TNF
GC
HDL
IFNβ
IL10
IL13
IL4
IL4+upLPS
upLPS+Immune complex (IC)
sLPS+IC
P3C
P3C+PGE2
PGE2
TNF+P3C
TNF+PGE2
TNF+PGE2+P3C (TPP)
TPP+IFNβ
TPP+IFNβ+IFNg
LA
LiA
OA
PA
SA
classical
intermediate
non-classical
C1Q+
CD1c+
CD141+
pDC
Monocyte
cDC
monocyte
C1Qhi
FOLR2+
Macrophage
C1qhi
CD1c +
CD141c +
moDC
Healthy Control
Active COVID
Convalescent COVID
* p<0.05 + p(adj) <0.1
Mean expression in group
0.0
0.5
1.0

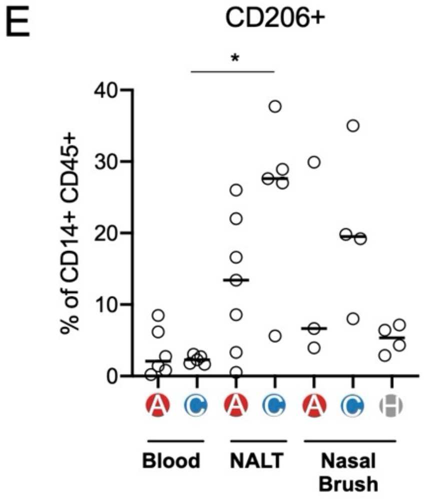
E
CD206+
*
% of CD14+ CD45+
0
10
20
30
40
Blood
NALT
Nasal Brush

**Figure S3.**

(A) Expression of canonical gene expression markers by assigned nasal MNP subset cell label. Assigned cell labels on y-axis, x-axis shows canonical marker genes grouped by cell type to which they correspond. Size of point indicates fraction of cells in each group expressing corresponding gene, colour of point indicates scaled mean expression of corresponding gene in each named cell type group.
(B) Additional images for figure 3F. Confocal immunofluorescence microscopy of NALT tissue from Convalescent COVID-19 subject showing spatial localisation of macrophage subsets relative to respiratory epithelium (EpCAM, white) and B cell follicles (CD19, yellow). FolRβ, green; CD163, red; CD206, blue.
(C) Expanded heatmap for figure 3G, indicating expression of genes associated with macrophage polarisation in all MNP subset cell types, split by disease type and sample type, calculated as scaled geneset expression scores (AddModuleScore [3]) using reference experimentally derived macrophage stimulation gene-sets [4]. Stimulation agent used for each gene-set shown on y-axis label, in correspondence to the original publication. H, Healthy control; A, Active COVID-19; C, Convalescent COVID-19.
(D) Comparative expression of macrophage stimulation and polarisation signatures shown in figure S3C [4], comparing in NALT monocyte, monocyte derived dendritic cell (moDC) and convention dendritic cell subsets, where MoDC have been segregated from conventional CD1c+ dendritic cells.
(E) Column scatter graph showing CD206 expression as a percentage of CD45+ CD14+ live single cells in blood (PBMC), NALT and inferior turbinate nasal brushing samples, by disease group. Each point represents an individual subject sample. A, Active COVID-19; C, Convalescent COVID-19, H, Healthy control. *, p(adj)<0.05 (Tukey HSD)

#### Supplementary 4

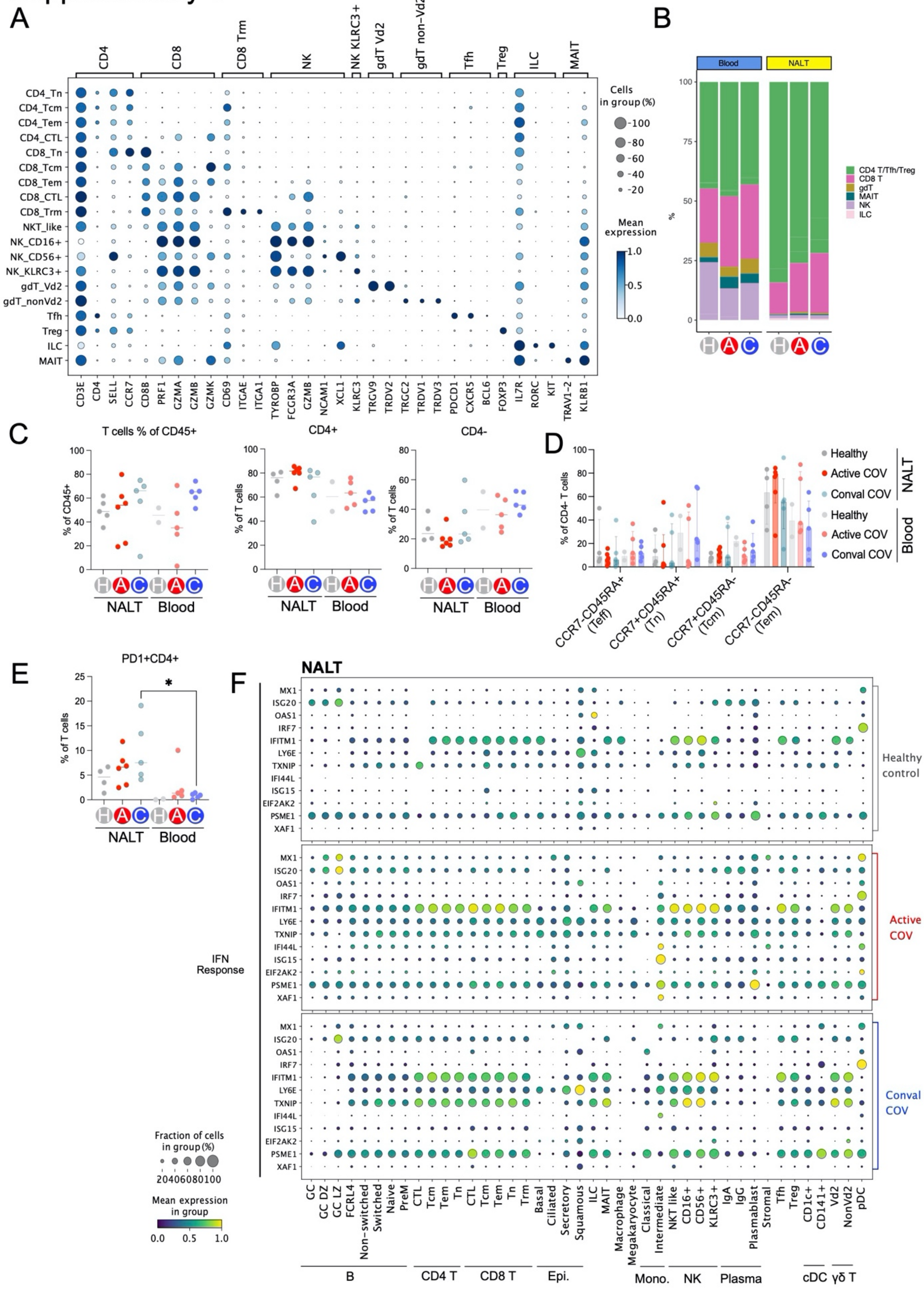

K

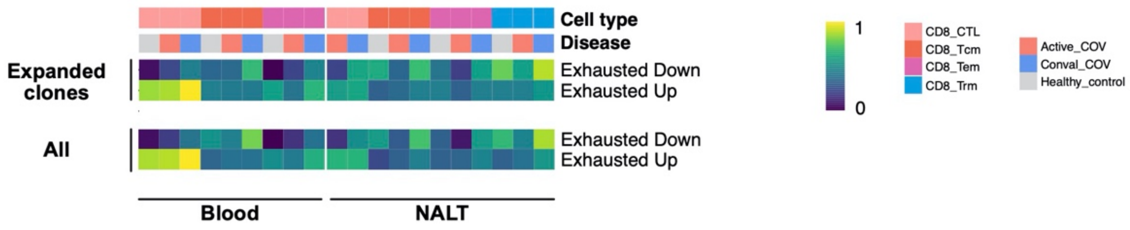
Cell type
Disease
Expanded clones
Exhausted Down
Exhausted Up
All
Exhausted Down
Exhausted Up
Blood
NALT
1
0
CD8_CTL
CD8_Tcm
CD8_Tem
CD8_Trm
Active_COV
Conval_COV
Healthy_control

L

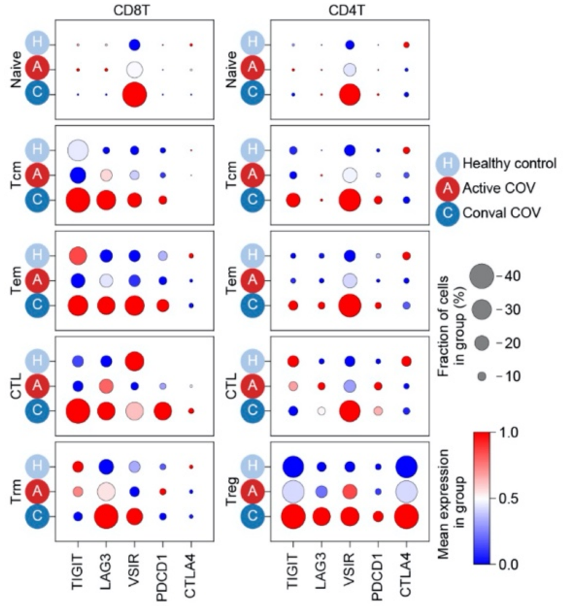
CD8T
CD4T
Naive
Tcm
Tem
CTL
Trm
Treg
TIGIT
LAG3
VSIR
PDCD1
CTLA4
Healthy control
Active COV
Conval COV
Fraction of cells in group (%)
40
30
20
10
Mean expression in group
1.0
0.5
0.0

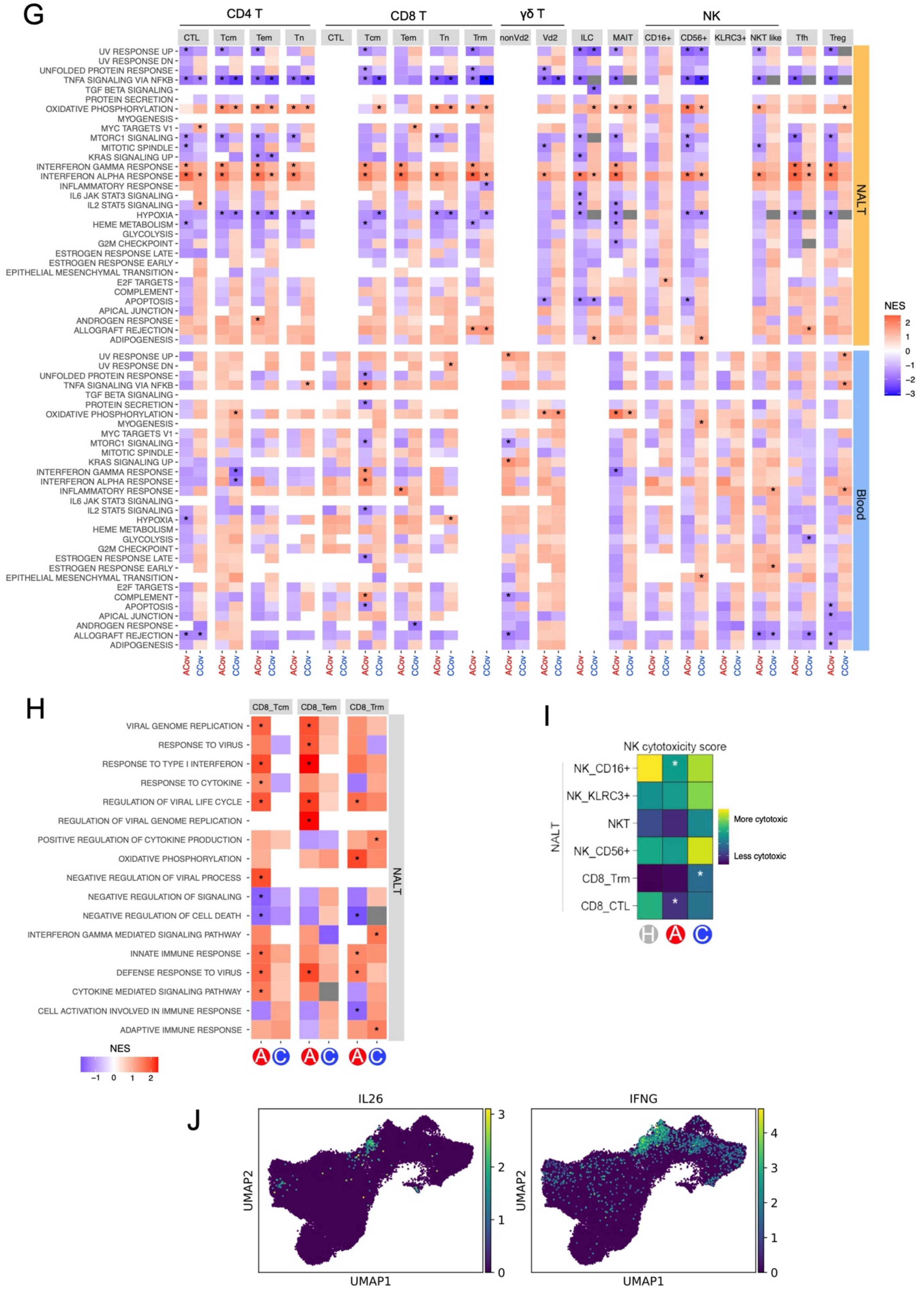
G
CD4 T
CD8 T
γδ T
NK
CTL
Tcm
Tem
Tn
Trm
nonVd2
Vd2
ILC
MAIT
CD16+
CD56+
KLRC3+
NKT like
Tfh
Treg
UV RESPONSE UP
UV RESPONSE DN
UNFOLDED PROTEIN RESPONSE
TNFA SIGNALING VIA NFKB
TGF BETA SIGNALING
PROTEIN SECRETION
OXIDATIVE PHOSPHORYLATION
MYOGENESIS
MYC TARGETS V1
MTORC1 SIGNALING
MITOTIC SPINDLE
KRAS SIGNALING UP
INTERFERON GAMMA RESPONSE
INTERFERON ALPHA RESPONSE
INFLAMMATORY RESPONSE
IL6 JAK STAT3 SIGNALING
IL2 STAT5 SIGNALING
HYPOXIA
HEME METABOLISM
GLYCOLYSIS
G2M CHECKPOINT
ESTROGEN RESPONSE LATE
ESTROGEN RESPONSE EARLY
EPITHELIAL MESENCHYMAL TRANSITION
E2F TARGETS
COMPLEMENT
APOPTOSIS
APICAL JUNCTION
ANDROGEN RESPONSE
ALLOGRAFT REJECTION
ADIPOGENESIS
NALT
Blood
NES
2
1
0
−1
−2
−3
ACov
CCov
H
CD8_Tcm
CD8_Tem
CD8_Trm
VIRAL GENOME REPLICATION
RESPONSE TO VIRUS
RESPONSE TO TYPE I INTERFERON
RESPONSE TO CYTOKINE
REGULATION OF VIRAL LIFE CYCLE
REGULATION OF VIRAL GENOME REPLICATION
POSITIVE REGULATION OF CYTOKINE PRODUCTION
OXIDATIVE PHOSPHORYLATION
NEGATIVE REGULATION OF VIRAL PROCESS
NEGATIVE REGULATION OF SIGNALING
NEGATIVE REGULATION OF CELL DEATH
INTERFERON GAMMA MEDIATED SIGNALING PATHWAY
INNATE IMMUNE RESPONSE
DEFENSE RESPONSE TO VIRUS
CYTOKINE MEDIATED SIGNALING PATHWAY
CELL ACTIVATION INVOLVED IN IMMUNE RESPONSE
ADAPTIVE IMMUNE RESPONSE
NALT
NES
−1
0
1
2
A
C
I
NK cytotoxicity score
NK_CD16+
NK_KLRC3+
NKT
NK_CD56+
CD8_Trm
CD8_CTL
NALT
More cytotoxic
Less cytotoxic
H
A
C
J
IL26
IFNG
UMAP1
UMAP2
0
1
2
3
4

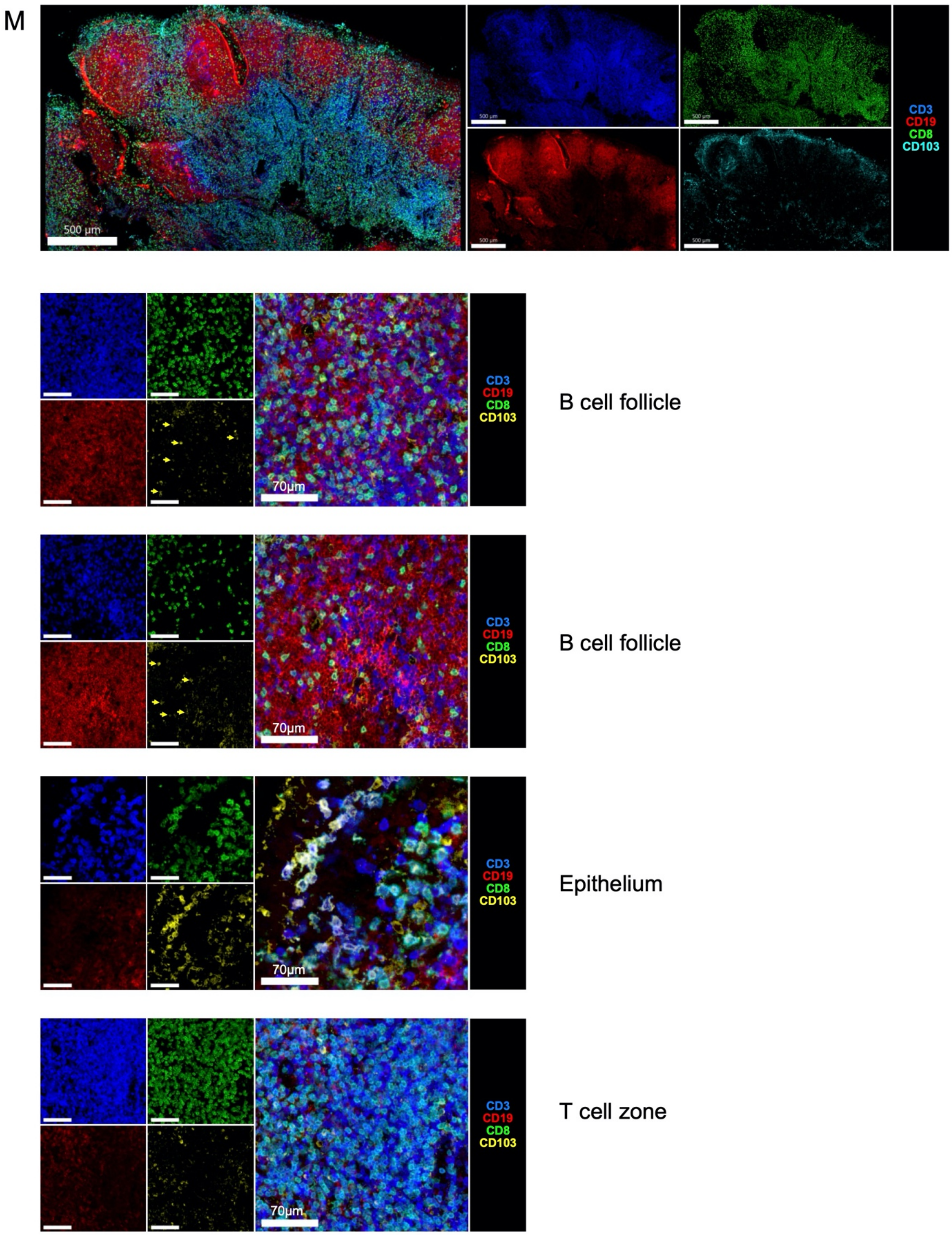
M
500 µm
CD3
CD19
CD8
CD103
70µm
B cell follicle
70µm
B cell follicle
70µm
Epithelium
70µm
T cell zone

#### Figure S4

(A) Expression of canonical gene expression markers by assigned T/innate lymphocyte subset cell label. Assigned cell labels on y-axis, x-axis shows canonical marker genes grouped by cell type to which they correspond. Size of point indicates fraction of cells in each group expressing corresponding gene, colour of point indicates scaled mean expression of corresponding gene in each named cell type group.

(B) Stacked bar charts showing combined proportional representation of CD4 T cells, CD8 T cells and innate lymphocytes, grouped into broad cell type categories and split by sample type (Blood/NALT) and by disease status. H, Healthy control; A, Active COVID-19; C, Convalescent COVID-19

(C) Grouped scatter plot of T cell proportions, as determined by flow cytometry, split by disease type and sample type. Left, CD3+ (T) cells as a proportion (%) of parent CD45+, live single cell gated population; Centre, CD4+ cells as a proportion of parent CD3+, CD45+, live single cell gated population; Right, CD4- cells as a proportion of parent CD3+, CD45+, live single cell gated population. Each point represents an individual subject sample. Lines represent group median value.

(D) Grouped scatter and bar plot of CD4- T (CD3+CD45+, live singlets) cell sub population proportions by CCR7 and CD45RA expression, as determined by flow cytometry. Each point represents an individual subject sample. Bars represent median values, with lines showing interquartile range. Teff, T effector; Tn, T naïve; Tcm, T central memory; Tem, T effector memory.

(E) Grouped scatter plot of PD1+CD4+ (Tfh) cells as a percentage of parent CD3+CD45+ live single cell population, as determined by flow cytometry. Each point represents an individual subject sample. Lines represent group median value. *, adjusted $p<0.05$ (Šidák multiple comparisons test). H, healthy control; A, Active COVID-19, C, Convalescent COVID-19.

(F) Scaled expression of selected leading edge genes in NALT cell types that were enriched in Hallmark Interferon Alpha Response and Hallmark Interferon Gamma Response (IFN Response) gene sets following gene set enrichment analysis of differentially expressed genes in COVID-19 subjects compared with healthy controls. Size of point indicates fraction of cells in each group expressing corresponding gene, colour of point indicates scaled mean expression of corresponding gene in each named cell type group. Cell types are shown on the x-axis, and genes on the y-axis. Epi, Epithelial; Mono., Monocyte.

(G) Extended heatmap from figure 4G, showing all significantly enriched Hallmark pathways in T/innate lymphocyte cell types following gene set enrichment analysis of differentially expressed genes between COVID-19 disease groups and healthy control subjects. Only pathways containing at least one significant enrichment ($p<0.05$ and Benjamini-Hochberg adjusted p value $<0.1$) in T cell types are shown. Colour indicates normalised enrichment score (NES), with red indicating greater pathway enrichment in disease group compared with healthy controls, and blue indicating increased pathway enrichment in healthy controls than disease group. Results are split by sample type (NALT/Blood)

(H) Selected significantly enriched Gene Ontology(GO) terms in NALT CD8 T cell memory subsets, following Geneset Enrichment Analysis of differentially expressed genes in COVID-19 disease states compared with healthy controls. NES, Normalised Enrichment Score, *, p value $<0.05$ + Benjamini-Hochberg adjusted p value $<0.1$. A, Active COVID-19; C, Convalescent COVID-19.

(I) Heatmap showing Gene-set scoring (AddModuleScore) of KEGG Natural killer cell mediated cytotoxicity gene set in NALT CD8 T and NK cell subsets. *, p value $<0.05$

(J) UMAP showing expression of IL26 and IFNG within subset re-clustered T/innate lymphocyte UMAP seen in figure 4A. Scale bars indicate level of gene expression.

(K) Extended heatmap from figure 4K, showing enrichment of Human HIV exhaustion signatures [5] split by cell type, sample type, disease type and clonal expansion (TCR clone size ≥2) status. Exhausted Up indicates genes upregulated in exhausted vs non-exhausted CD8 T cells, and Exhausted Down indicated genes downregulated in exhausted vs non-exhausted CD8 T cells. Top two rows indicate cell type and disease status for each column, corresponding with the legend (right)

(L) Dotplot showing expression of canonical T cell exhaustion marker genes in CD4 and CD8 T cell subsets. Size of point indicates fraction of cells in each group expressing corresponding gene, colour of point indicates scaled mean expression of corresponding gene in each named cell type group. H, Healthy control, A, Active COVID-19, C, Convalescent COVID-19

(M) Extended imaging from Figures 4E-F, showing confocal immunofluorescence microscopy image of NALT tissue in a Convalescent COVID-19 subject showing CD8 Trm (CD3+CD8+CD103+) spatial localisation at lower magnification (top), with high magnification images seen below from the B cell follicle, epithelium and T cell zone, showing CD8 and CD103 co-localisation (indicated with arrows in the B cell follicle images to aid identification).

### Supplementary Figure 5

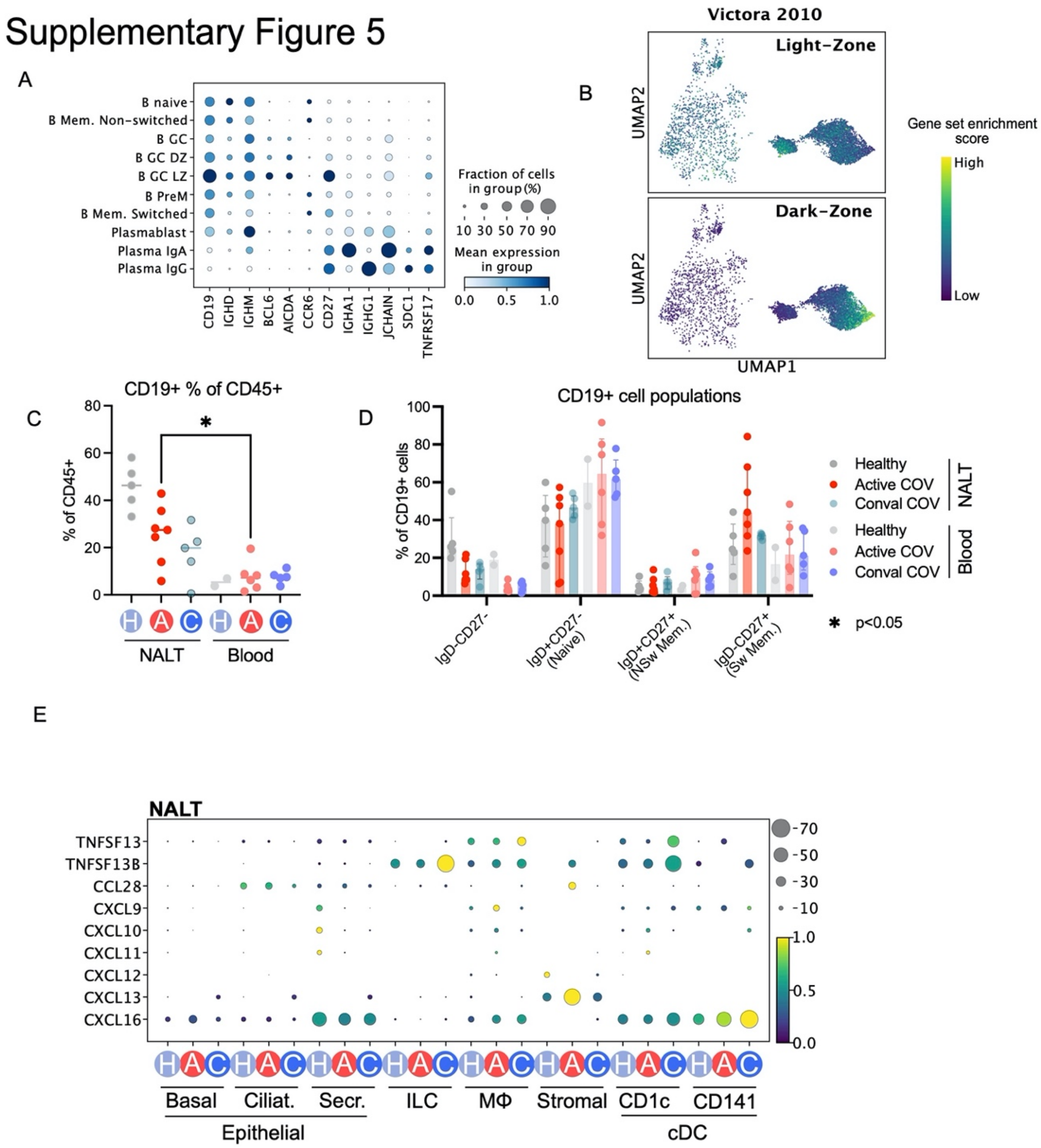

F

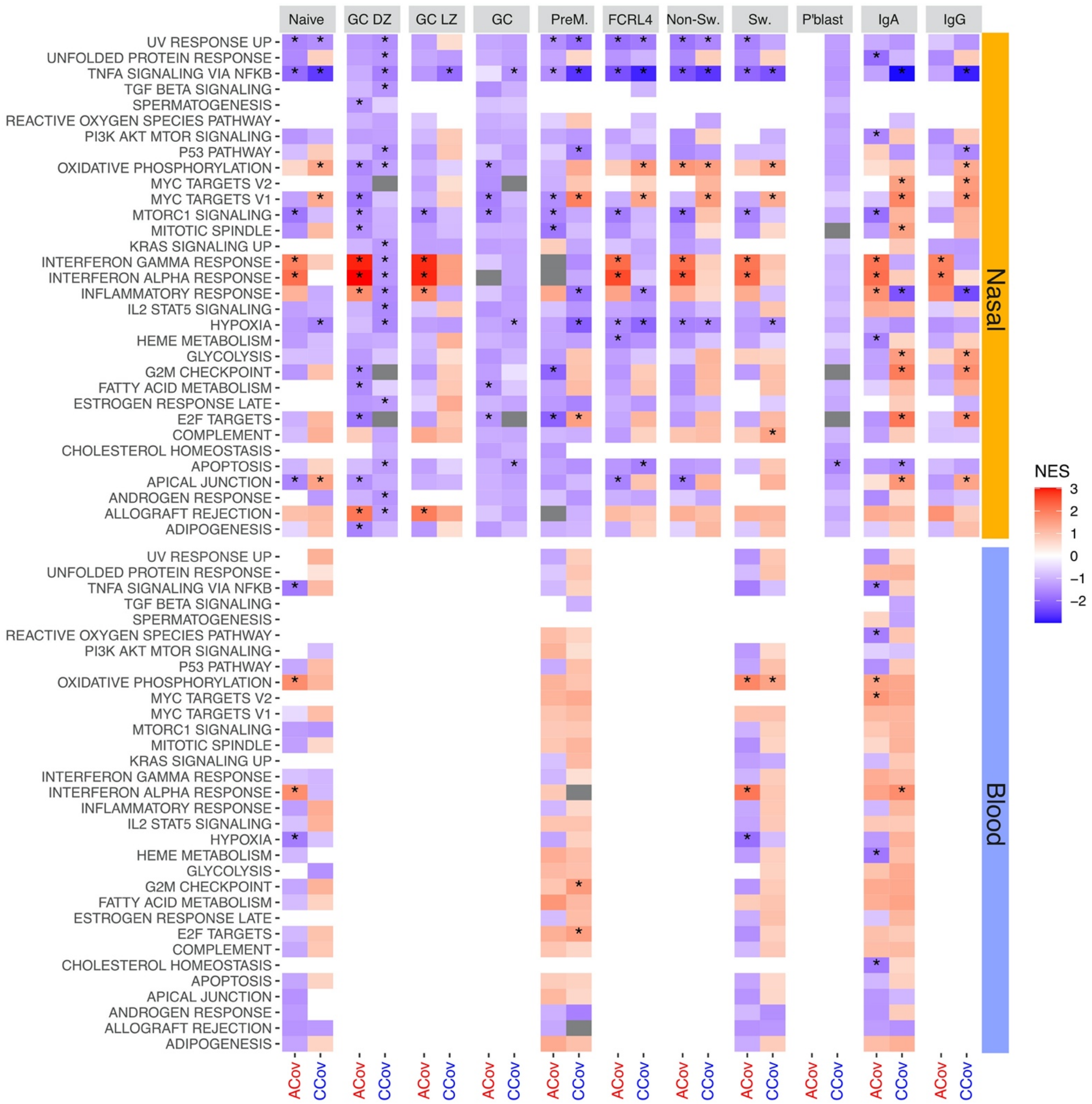
Naive
GC DZ
GC LZ
GC
PreM.
FCRL4
Non-Sw.
Sw.
P'blast
IgA
IgG
UV RESPONSE UP
UNFOLDED PROTEIN RESPONSE
TNFA SIGNALING VIA NFKB
TGF BETA SIGNALING
SPERMATOGENESIS
REACTIVE OXYGEN SPECIES PATHWAY
PI3K AKT MTOR SIGNALING
P53 PATHWAY
OXIDATIVE PHOSPHORYLATION
MYC TARGETS V2
MYC TARGETS V1
MTORC1 SIGNALING
MITOTIC SPINDLE
KRAS SIGNALING UP
INTERFERON GAMMA RESPONSE
INTERFERON ALPHA RESPONSE
INFLAMMATORY RESPONSE
IL2 STAT5 SIGNALING
HYPOXIA
HEME METABOLISM
GLYCOLYSIS
G2M CHECKPOINT
FATTY ACID METABOLISM
ESTROGEN RESPONSE LATE
E2F TARGETS
COMPLEMENT
CHOLESTEROL HOMEOSTASIS
APOPTOSIS
APICAL JUNCTION
ANDROGEN RESPONSE
ALLOGRAFT REJECTION
ADIPOGENESIS
Nasal
Blood
NES
3
2
1
0
−1
−2
ACov
CCov

G

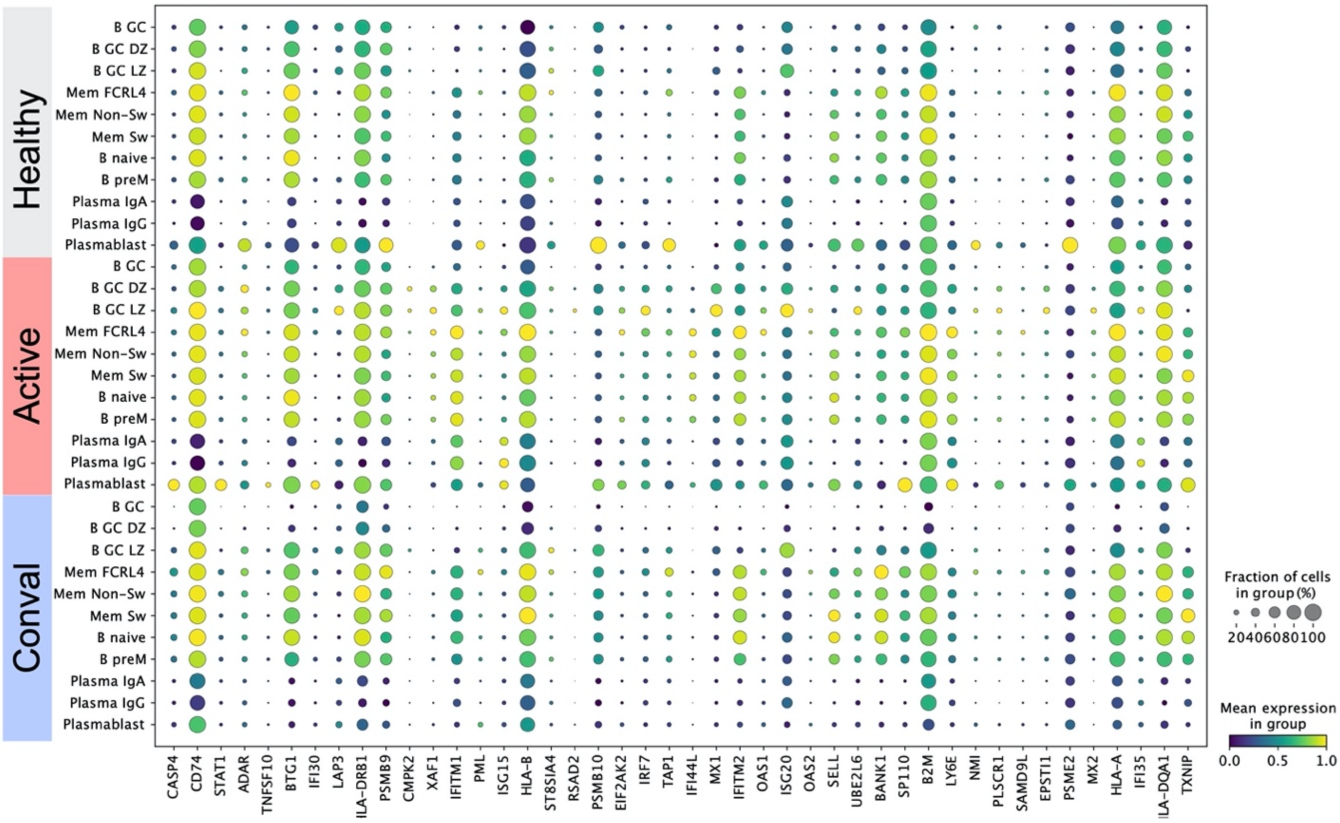

Healthy
Active
Conval
B GC
B GC DZ
B GC LZ
Mem FCRL4
Mem Non-Sw
Mem Sw
B naive
B preM
Plasma IgA
Plasma IgG
Plasmablast
CASP4
CD74
STAT1
ADAR
TNFSF10
BTG1
IFI30
LAP3
ILA-DRB1
PSMB9
CMPK2
XAF1
IFITM1
PML
ISG15
HLA-B
ST8SIA4
RSAD2
PSMB10
EIF2AK2
IRF7
TAP1
IFI44L
MX1
IFITM2
OAS1
ISG20
OAS2
SELL
UBE2L6
BANK1
SP110
B2M
LY6E
NMI
PLSCR1
SAMD9L
EPSTI1
PSME2
MX2
HLA-A
IFI35
ILA-DQA1
TXNIP
Fraction of cells in group (%)
20406080100
Mean expression in group
0.0
0.5
1.0


H

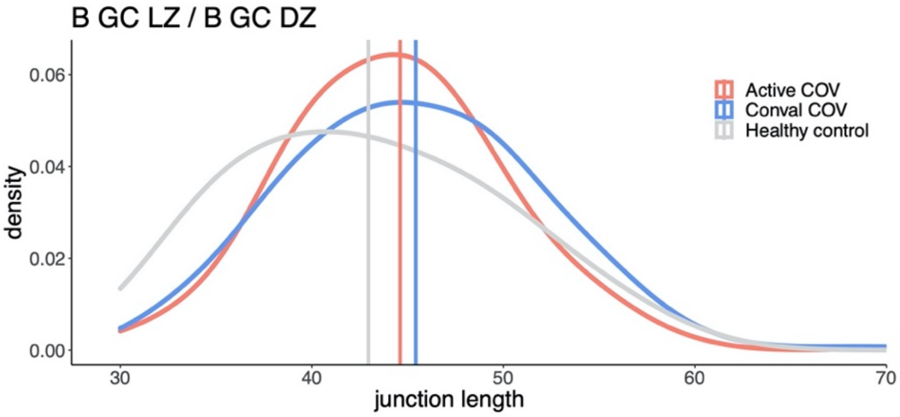

B GC LZ / B GC DZ
0.06
0.04
0.02
0.00
density
30
40
50
60
70
junction length
Active COV
Conval COV
Healthy control

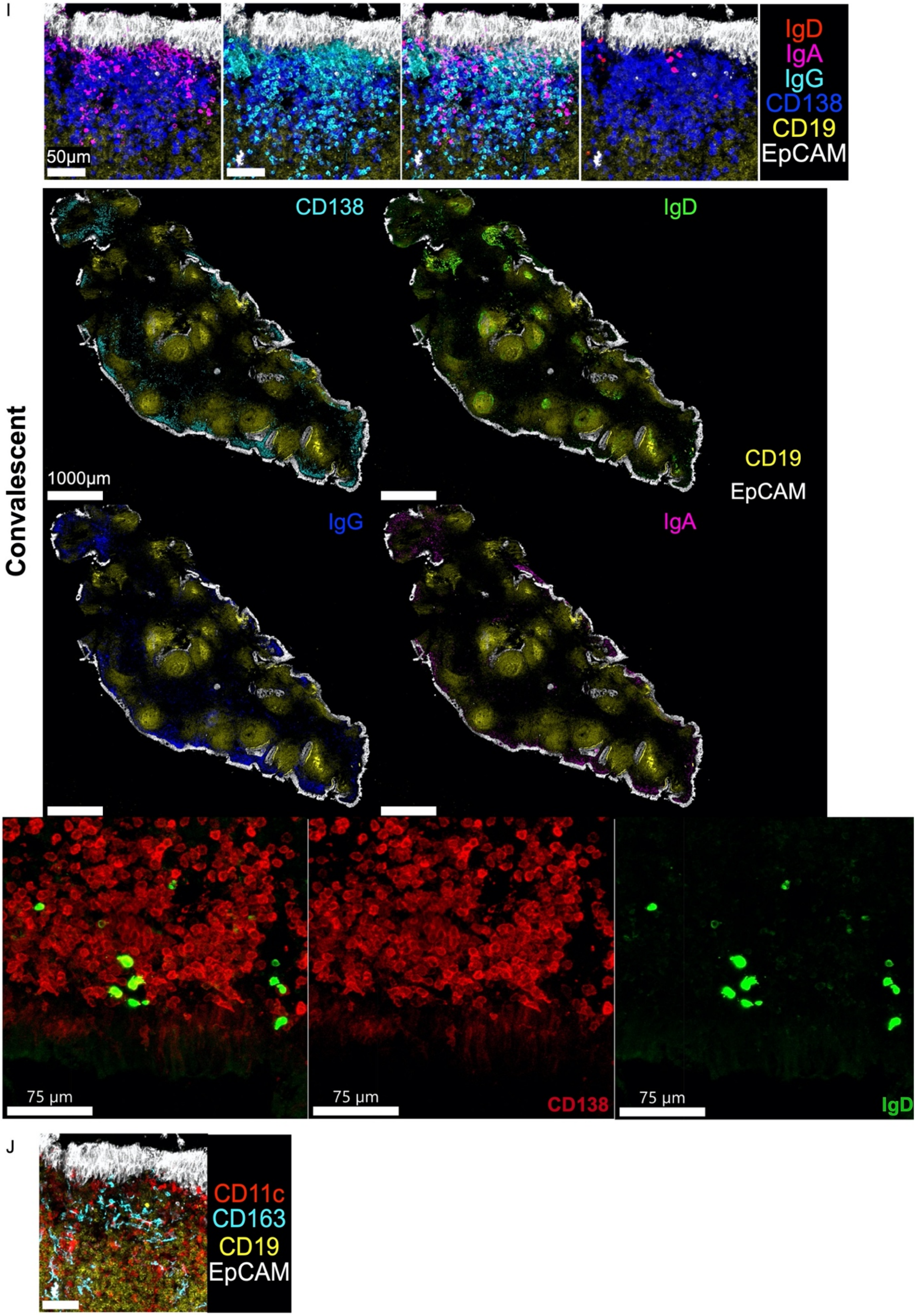
I
IgD
IgA
IgG
CD138
CD19
EpCAM
50µm
Convalescent
CD138
IgD
CD19
EpCAM
IgG
IgA
1000µm
75 µm
75 µm
CD138
75 µm
IgD
J
CD11c
CD163
CD19
EpCAM

#### Figure S5

(A) Expression of canonical gene expression markers by assigned B lymphocyte subset cell label. Assigned cell labels on y-axis, x-axis shows canonical marker genes grouped by cell type to which they correspond. Size of point indicates fraction of cells in each group expressing corresponding gene, colour of point indicates scaled mean expression of corresponding gene in each named cell type group.
(B) UMAP showing gene set enrichment score (AddModuleScore) of experimentally derived Germinal centre B cell signatures [6] in germinal centre B cell clusters.
(C) Grouped scatter plot of CD19+ (B) cells as a percentage of parent CD45+ live single cell population, as determined by flow cytometry. Each point represents an individual subject sample. Lines show group median value. Data is divided by disease and sample type. *, $p<0.05$ (Wilcoxon test) H, healthy control; A, Active COVID-19, C, Convalescent COVID-19.
(D) Grouped scatter and bar plot of CD19+ (B) cell sub population proportions by IgD and CD27 expression, as determined by flow cytometry. Each point represents an individual subject sample. Bars represent median values, with lines showing interquartile range. NSw Mem, Non-switched memory; Sw. Mem, Switched memory.
(E) Dotplot showing selected chemokine expression across NALT Epithelial cells, ILCs, Macrophages (MΦ), Stromal cells and conventional dendritic cells. Size of point indicates fraction of cells in each group expressing corresponding gene, colour of point indicates scaled mean expression of corresponding gene in each named cell type group.
(F) Extended heatmap from figure 5I, showing all significantly enriched Hallmark pathways in B lymphocyte/plasma cell types following gene set enrichment analysis of differentially expressed genes between COVID-19 disease groups and healthy control subjects. Only pathways containing at least one significant enrichment ($p<0.05$ and Benjamini-Hochberg adjusted p value $<0.1$) in B cell types are shown. Colour indicates normalised enrichment score (NES), with red indicating greater pathway enrichment in disease group compared with healthy controls, and blue indicating increased pathway enrichment in healthy controls than disease group. Results are split by sample type (NALT/Blood)
(G) Scaled dotplot of expression in B and plasma cell subsets of hallmark Interferon alpha and gamma response nasal leading edge genes from GSEA of differentially expressed genes in COVID-19 groups compared with healthy controls. Size of point indicates fraction of cells in each group expressing corresponding gene, colour of point indicates scaled mean expression of corresponding gene in each named cell type group.
(H) Density plot of BCR junction length in light zone and dark zone germinal centre B cells, split by condition. Vertical lines indicate mean junction length for each disease group. Active COVID-19 (Red), Convalescence COVID-19 (Blue), Healthy Control (Grey).
(I) Extended images from figure 5D. Confocal immunofluorescence microscopy showing localisation and isotype expression of plasma cells in NALT tissue from a convalescent COVID-19 subject. Top: high magnification images showing plasma cells (CD138, purple) and isotype expression in the sub-eptheilal region (IgD, Red; IgA, Pink; IgG, Cyan; CD138, Purple; CD19, Yellow; EpCAM, White). Middle: low magnification images of NALT tissue showing localisation of plasma cells and isotype expression through the NALT tissue section (CD138, Cyan; IgD, Green; IgG, Purple; IgA, Pink). Bottom: High magnification image showing IgD+ plasma cells (CD138, Red; IgD, Green).
(J) Extended image from figure 5H, showing figure 5H without CD138 staining.

### Supplementary 6

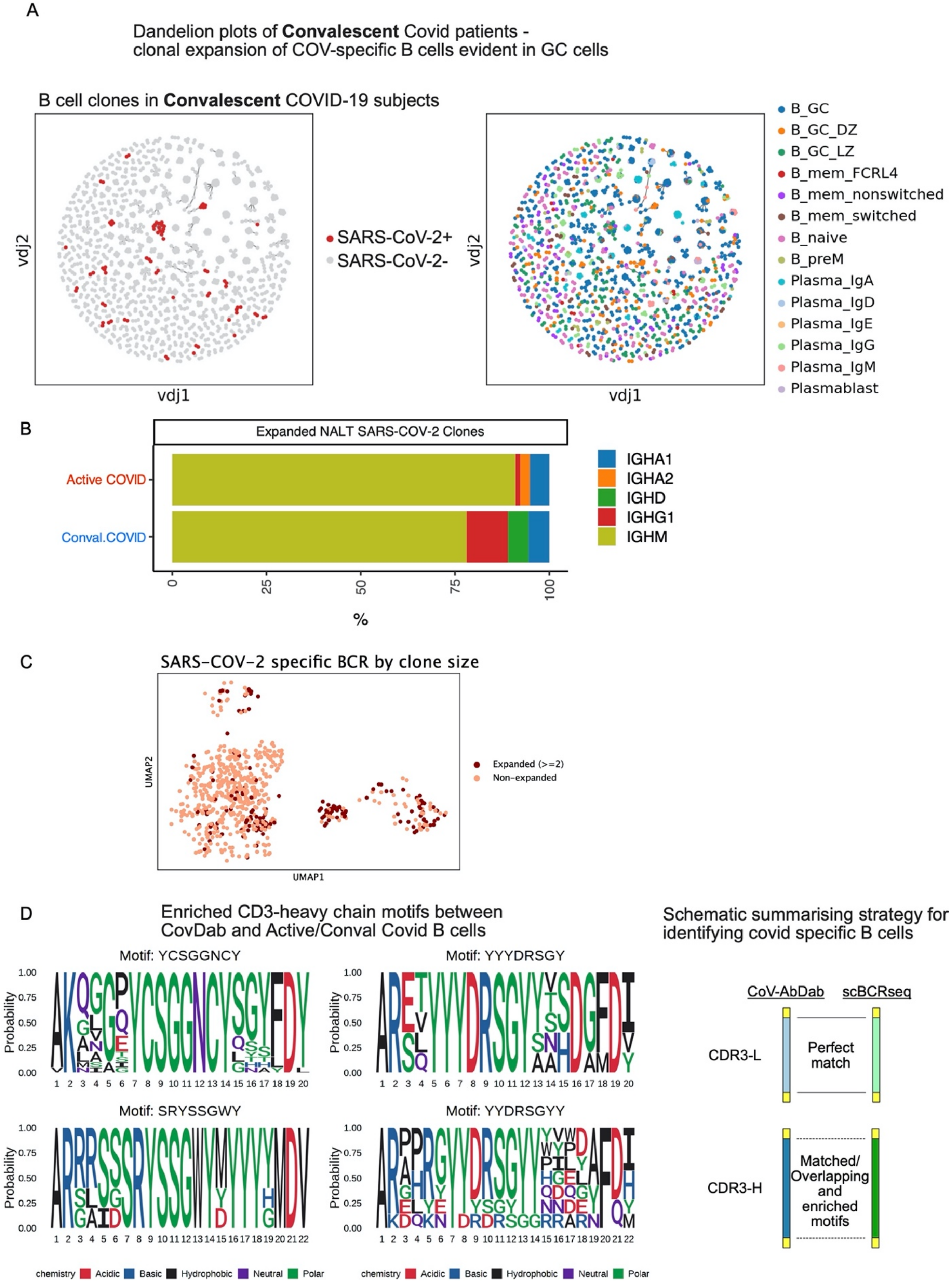

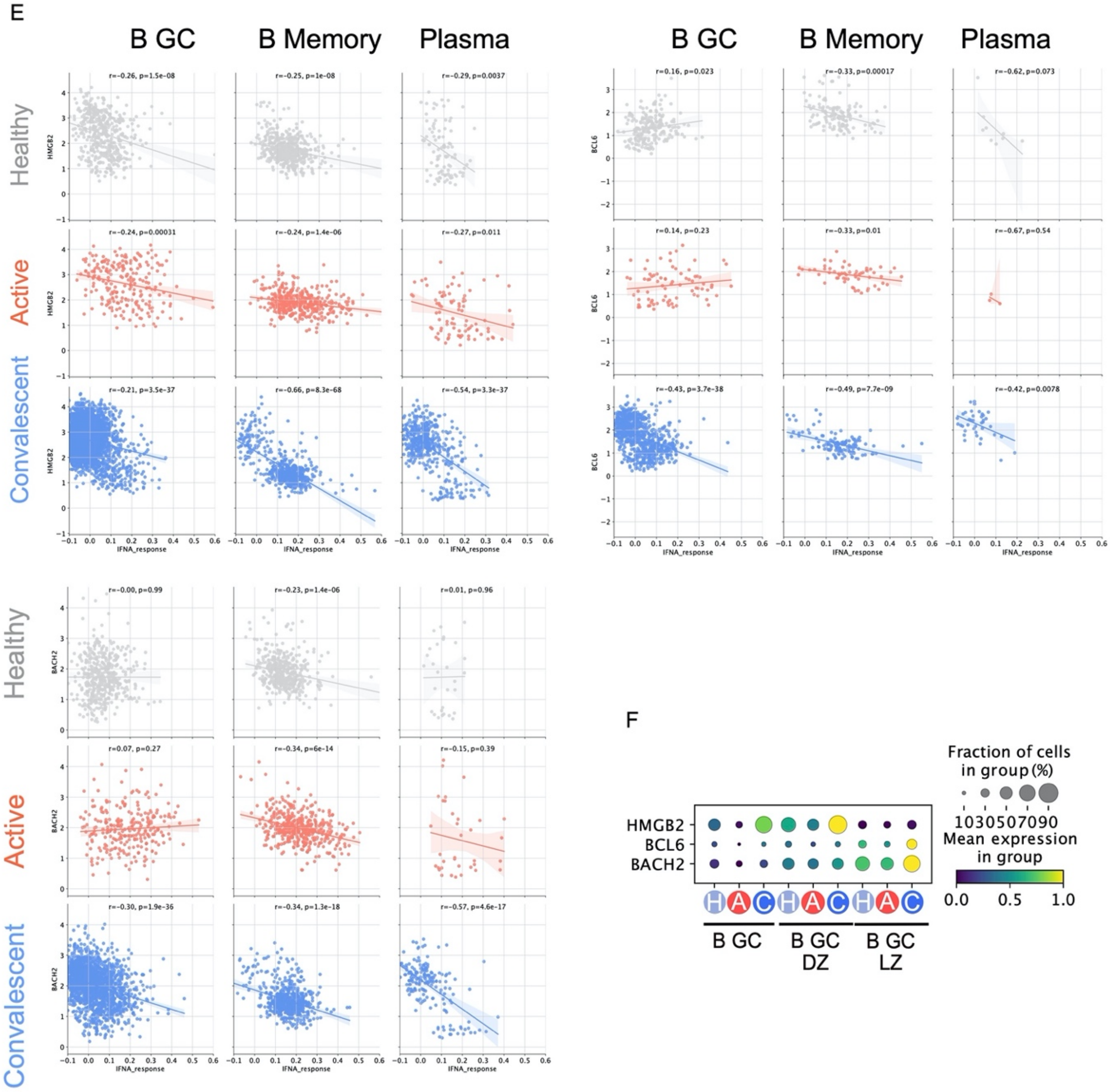
E
B GC
B Memory
Plasma
Healthy
Active
Convalescent
HMGB2
BCL6
BACH2
IFNA_response
r=-0.26, p=1.5e-08
r=-0.25, p=1e-08
r=-0.29, p=0.0037
r=-0.24, p=0.00031
r=-0.24, p=1.4e-06
r=-0.27, p=0.011
r=-0.21, p=3.5e-37
r=-0.66, p=8.3e-68
r=-0.54, p=3.3e-37
r=0.16, p=0.023
r=-0.33, p=0.00017
r=-0.62, p=0.073
r=0.14, p=0.23
r=-0.33, p=0.01
r=-0.67, p=0.54
r=-0.43, p=3.7e-38
r=-0.49, p=7.7e-09
r=-0.42, p=0.0078
r=-0.00, p=0.99
r=-0.23, p=1.4e-06
r=0.01, p=0.96
r=0.07, p=0.27
r=-0.34, p=6e-14
r=-0.15, p=0.39
r=-0.30, p=1.9e-36
r=-0.34, p=1.3e-18
r=-0.57, p=4.6e-17
F
Fraction of cells
in group (%)
1030507090
Mean expression
in group
0.0 0.5 1.0
HMGB2
BCL6
BACH2
H A C H A C H A C
B GC
B GC
DZ
B GC
LZ

**Figure S6**

(A) Single-cell BCR network plots for Convalescent COVID-19 subjects. Each circle/node corresponds to a single B cell with a corresponding set of BCR(s). Each clonotype is presented as a minimally connected graph with edge widths scaled to 1/d + 1 for edge weight d where d corresponds to the total (Levenshtein) edit distance of BCRs between two cells. Left-hand plot shows SARS-COV-2 specific clones (red) in the context of all B cell clones. Right-hand plot shows assigned cell type labels derived from gene expression data.
(B) Stacked bar plot showing proportional split by isotype of expanded (clone size >=2) SARS-COV-2 specific B cell clones by disease state.
(C) UMAP showing SARS-COV-2 specific B cell clones plotted using co-ordinates from B/Plasma UMAP seen in figure 5A, showing localisation of expanded SARS-COV-2 clones to the light zone germinal centre cluster. Cells forming part of expanded BCR clones (clone size >=2) are shown in dark red, with clonally unexpanded SARS-COV-2 specific B cells shown in pink.
(D) Left, Enriched CD3-heavy chain motifs between Cov-AbDab and B cells from Active/Conval COVID-19 subjects. Right, Schematic summarising strategy for identifying SARS-COV-2 specific B cells from the single cell CDR3-L and CDR3-H sequences shared with CoV-AbDab.
(E) Extended figures from figure 6K, showing plots split by cell type and disease type for interferon alpha response scores plotted against expression of HMGB2 (upper left), BCL6 (upper right), and BACH2 (lower left). Linear regression line is shown with a shaded area representing the 95% confidence interval. r, Pearson correlation statistic; p, p-value. *, $p<0.05$; **, $p<0.01$; ***, $p<0.001$; ns, $p>0.05$.
(F) Dotplot showing expression of genes shown in figure 6K in NALT germinal centre B cells, split by cell type and disease type. Size of point indicates fraction of cells in each group expressing corresponding gene, colour of point indicates scaled mean expression of corresponding gene in each cell type and disease group.
